# Etiology and epidemiology of branch dieback and fruit blight and necrosis of English walnut in France

**DOI:** 10.1101/2025.02.20.639244

**Authors:** Marie Belair, Adeline Picot, Cyrielle Masson, Marie-Neige Hébrard, Marie Debled, Aude Moronvalle, Benjamin Richard, Sylvie Tréguer, Léa Morvant, Amandine Henri-Sanvoisin, Yohana Laloum, Gaétan Le Floch, Flora Pensec

## Abstract

English walnut (*Juglans regia* L.) is an economically-important fruit crop worldwide, particularly in France, where it is the second most important fruit crop after apples in terms of cultivated area. Walnut orchards are targeted by numerous diseases, but new symptoms have been widely observed since 2015 in France, consisting in typical branch dieback and fruit blight and necrosis. Herein, both symptomatic and asymptomatic twigs and husks were collected from 12 commercial French walnut orchards from the two main production areas located in the South of France from 2020 to 2022. Overall, Botryosphaeriaceae, *Colletotrichum, Diaporthe*, and *Fusarium* species were consistently isolated from symptomatic husks and twigs. Among these taxa, *B. dothidea* and *C. godetiae* were significantly more associated with symptomatic husks, contrary to *D. eres* and *F. juglandicola,* more frequently isolated from symptomatic twigs, as also confirmed by ITS2 metabarcoding sequencing. Tissue type was the primary factor significantly impacting the composition of the fungal communities (R² = 16.6%), followed by sampling campaign and production area (R² = 11.6% and 4.6%, respectively). We also found that *Neofusicoccum* and *Diaporthe* were significantly more associated with warm and dry years, unlike *Colletotrichum*. Pathogenicity tests revealed that only Botryosphaeriaceae and *Diaporthe* species were pathogenic on walnut fruits and twigs, with *B. dothidea* and *N. parvum* being the most aggressive species. Several significant positive and negative correlations were identified based on SparCC co-occurrence networks and correlograms based on prevalence data. In particular, the positive correlation between *Colletotrichum* and *Fusarium* species on symptomatic twigs was highlighted by both methods. This work is the first comprehensive etiological and epidemiological study of branch dieback and fruit blight and necrosis of English walnut in France.

**Author Summary:** English walnut is an economically important crop, mainly produced in China, in the United States and in Turkey while in Europe, France is one of the biggest producers. Although widespread in nut orchards abroad, new symptoms, including fruit necroses and branch dieback, were observed for the first time in 2015 in France. We then conducted sampling campaigns between 2020 and 2022 to collect walnut husks and twigs in the two main French production areas. Exploration of the associated microbiota revealed the presence of six predominant fungal species, even in samples without symptoms. We then investigated the capacity of these species to cause lesions on twigs and fruits. Among all species, *Neofusicoccum parvum* was the most aggressive on both organs while other species were mainly aggressive on only twigs or husks. In addition, we evidenced that the presence of some species was favored by warm and dry weather or inversely cold and wet weather. We also found negative correlations between certain species (in particular between *N. parvum* and *Diaporthe eres*), suggesting that some species probably prevented the other from further colonization. Overall, we identified the species specifically associated with the disease in France and the factors influencing the disease risk. This work will enable us to better identify the best strategies to combat this disease.

## Introduction

English walnut (*Juglans regia* L.), or common walnut, is the most cultivated species of the *Juglans* genus for its fruit production [1]. Worldwide production of walnuts is estimated to reach approximately 3.5 million tons, of which 37,000 tons are produced in France [2]. The French walnut industry is mainly concentrated in the south of France and is divided into two main production areas: the South-East area (Auvergne-Rhône-Alpes region) and the South-West area (Nouvelle-Aquitaine and Occitanie regions). Over the last thirty years, the French walnut production area has increased by 148.8%, showing an increasing interest of the producers in walnut cultivation [2]. However, walnut growers have to face many challenges, including the management of pests and diseases, resulting in severe yield losses. Among them, the bacterial blight caused by *Xanthomonas campestris* pv. *juglandis* and the anthracnose caused by *Ophiognomonia leptostyla* have been historically encountered [3,4]. These challenges were further compounded by the development of a form of anthracnose caused by species of the genus *Colletotrichum* and responsible for fruit losses of up to 70%, since 2007 onwards [5]. More recently, in 2015, emergent symptoms, rarely observed so far in France, were reported at a nationwide level [6]. These new symptoms were characterized by fruit necrosis and blight, twig and shoot necrosis further leading to branch and even tree dieback associated with defoliation and dead buds, as well as branch canker [6].

In a previous survey conducted in the South-West of France in 2019, fungi isolated from symptomatic walnut husks and twigs showing the latter symptoms were identified as mainly belonging to the Botryosphaeriaceae family, including *Botryosphaeria* spp., *Dothiorella* spp. and *Neofusicoccum* spp., and the *Diaporthe* genus (teleomorph of *Phomopsis* spp.), as well as to the *Colletotrichum* and *Fusarium* genera to a lesser extent [7]. The symptomatology combined with the identification of species associated with symptoms suggested that the “*Botryosphaeria* and *Phomopsis* canker and blight disease”, also known as “Branch dieback and shoot blight disease”, was present in French walnut orchards, in association with other phytopathogenic species. This disease is known to be widespread in California since the early 2000s [8], and has more recently been reported in European countries such as the Czech Republic [9], Spain [10], Italy [11], Romania [12], and Hungary [13]. These studies reported up to ten species of the Botryosphaeriaceae family and up to two *Diaporthe* species, isolated from samples showing typical dieback symptoms of English walnut. Moreover, other species such as *Alternaria alternata, Cadophora* spp., *Colletotrichum acutatum, Cytospora* spp., *Fusarium* spp., and *Phaeoacremonium* spp. were identified, but to a lesser extent. Overall, these studies highlighted the frequent co-occurrence of species of Botryosphaeriaceae and *Diaporthe* on symptomatic walnut tissues.

Members of the Botryosphaeriaceae family and *Diaporthe* genus are considered to be endophytes, saprophytes and pathogens. Their ability to develop in tissues without developing symptoms, leading to latent infection, and to switch to a pathogenic lifestyle is widely acknowledged [14,15]. These species are responsible for numerous diseases on a wide range of woody and herbaceous plants, such as grapevine [16], pistachio [17] and oak [18,19], and are frequently found in co-occurrence, prompting the evaluation of their aggressiveness in single or mixed inoculations [20,21]. Overall, *B. dothidea* and *Diaporthe* spp. were more often associated with weak disease severity on walnut trees, whereas *Lasiodiplodia* and *Neofusicoccum* species were frequently associated with a higher aggressiveness [8–11,22–24].

Botryopshaeriaceae and *Diaporthe* were also often reported in association with secondary genera, including *Colletotrichum* and *Fusarium*. The former genus is composed of worldwide distributed pathogenic species responsible for numerous diseases [25], including anthracnose on walnut trees. Moreover, it was also identified in walnut fruits showing brown apical necrosis symptoms [26,27], as well as in apparently healthy walnut leaves [28]. As for the *Fusarium* genus, it is composed of species considered as endophytes, saprophytes and pathogens. Some species have already been identified as endophytes of apparently healthy walnut tissues [26,28,29], but many of them were also collected from symptomatic tissues and associated with walnut diseases. This mainly includes the brown apical necrosis (BAN) disease [26,30] and twig and stem canker symptoms, mainly associated with *F. solani* [31–34].

The emerging and widespread nature of these symptoms urged us to carry out an etiological and epidemiological study of branch dieback and fruit necrosis of English walnut on a larger scale in France. Furthermore, given the complexity of these diseases where several pathogenic species are often co-occurring, we also focused on the whole pathobiome, taking into account potential interactions. To do so, the objectives of this work were (i) to isolate and identify the main fungal species associated with branch dieback and fruit necrosis of English walnut based on morphological and molecular characterization, and evaluate their pathogenicity on walnut twigs and fruits in single inoculations, (ii) to describe more extensively the fungal pathobiome composition based on a metabarcoding approach, (iii) to study the climatic factors influencing the pathobiome based on culture-dependent and -independent approaches, and (iv) to determine potential interactions between pathogenic species and with members of the phytomicrobiome based on co-occurrence networks and correlation analysis.

## Results

### Field surveys, fungal isolation and identification

Samples with typical dieback symptoms were collected during the four sampling campaigns between 2020 and 2022 and included blighted and necrosed fruits (mummified in some cases), and defoliation and necroses on shoots and twigs as well as cankers with internal discoloration (Fig 1). Fungal cultures were obtained from all the 476 symptomatic samples (188 walnut husks and 288 twigs), as well as 95 out of 105 asymptomatic samples (33 walnut husks and 72 twigs), yielded to a collection of 1,873 fungal isolates excluding isolates from the same species collected from the same tissue. A total of 95% of isolates collected from symptomatic samples were represented by six endophytic/pathogenic fungal genera and the Didymellaceae family. Based on macroscopic observations and ITS sequencing, *Diaporthe* was the most collected genus, representing 26.2% of the isolates, followed by *Fusarium* (24.5%) and *Colletotrichum* (16.0%). The isolation of members of Botryosphaeriaceae, represented by *Botryosphaeria* (8.9%) and *Neofusicoccum* (9%), was less significant. Finally, Didymellaceae and *Alternaria* represented 5.5% and 4.5% of the fungal collection, respectively (**Fig 2A**). The same seven taxa were also found predominant in asymptomatic samples (309 isolates out of 325) with *Diaporthe* (32% of total isolates from asymptomatic samples), followed by the Didymellaceae family (16.6%), *Colletotrichum* (14.5%)*, Alternaria* (11.7%) and *Fusarium* (9.8%). Finally, the two Botryosphaeriaceae genera were represented by only 10.5% of the collected isolates (**Fig 2B)**.

**Fig 1.**
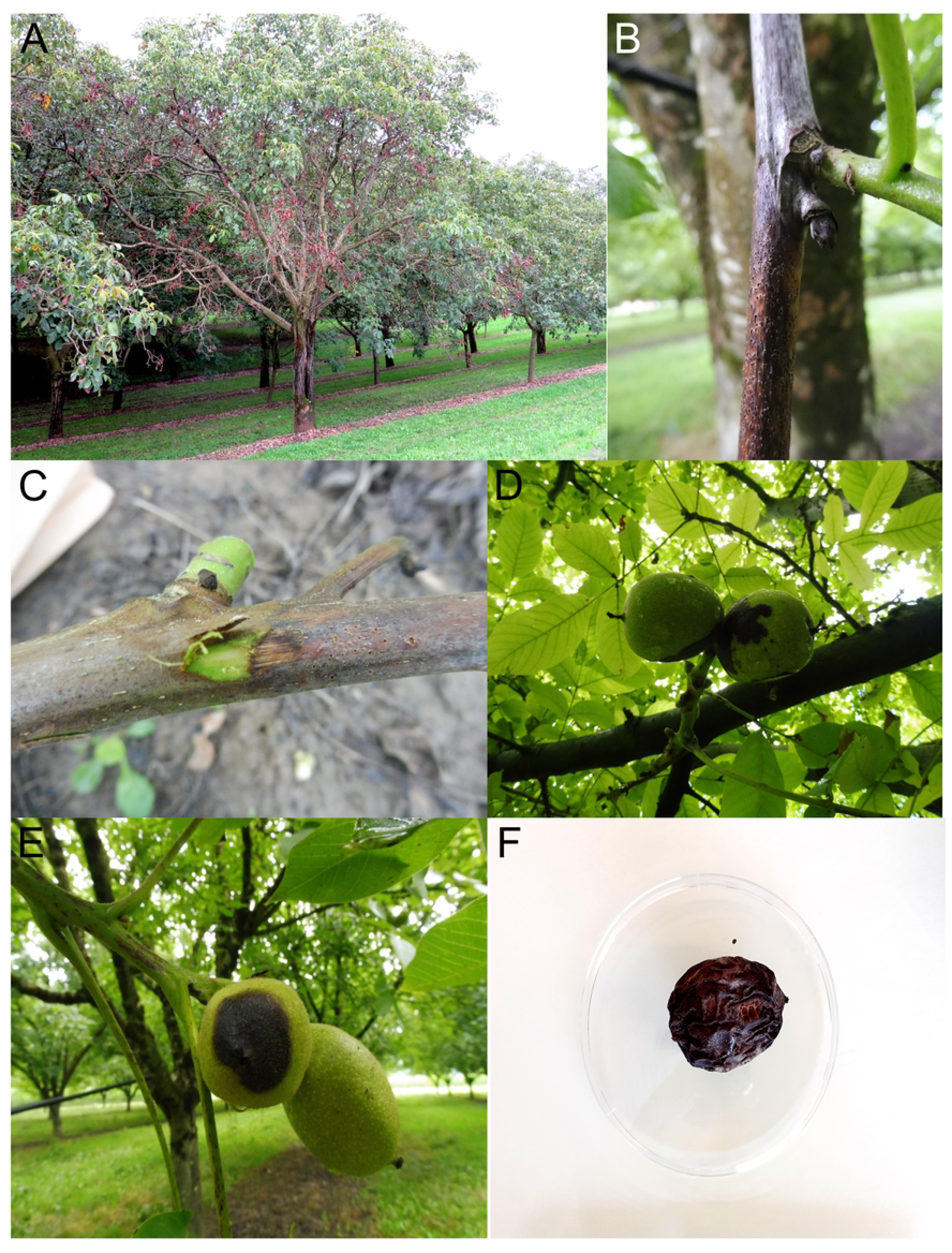
Symptoms of branch dieback and fruit blight and necrosis of English walnut in southern France. (A) Walnut tree showing decayed branches and necrotic foliage. (B) Typical symptoms of twig blight on young twig and dead bud. (C) Wood discoloration of blighted twig. (D & E) Typical fruit necrosis symptoms. (F) Mummified fruit.

**Fig 2.**
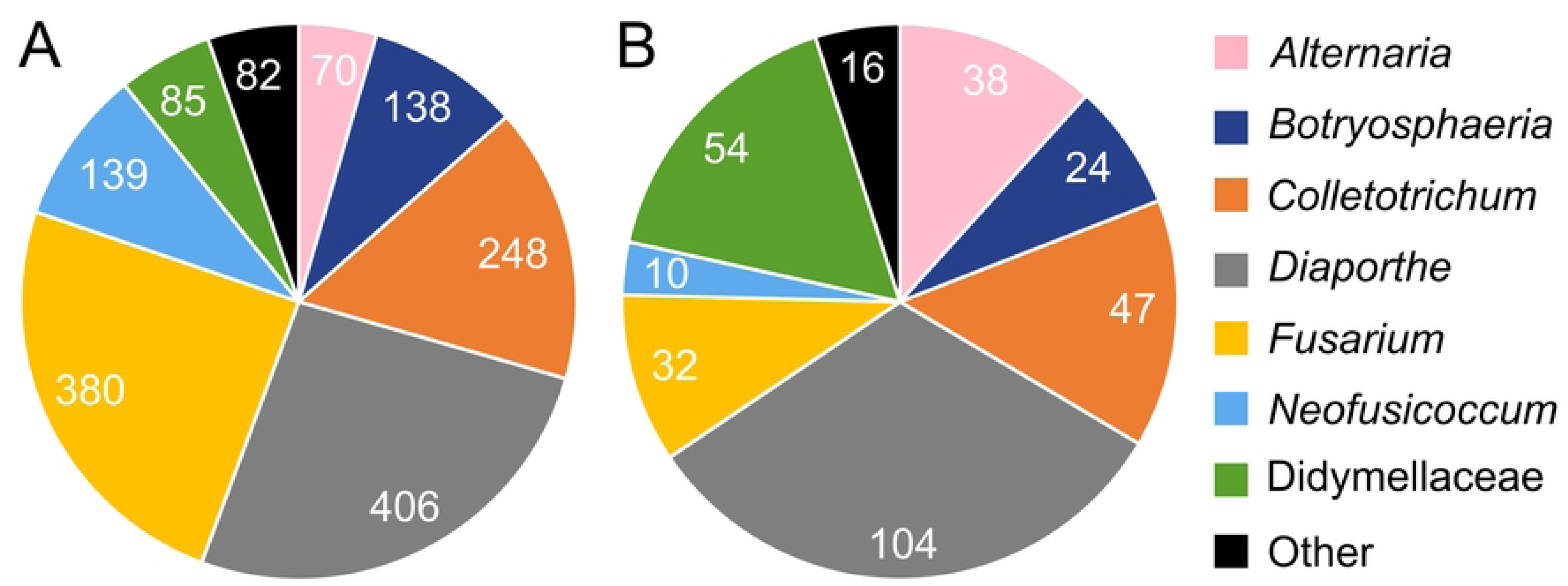
Distribution of species collected from symptomatic (A) and asymptomatic (B) husks and twigs over the four sampling campaigns. Values indicates the total number of isolates assigned to each taxa.

As ITS sequencing is insufficient to provide accurate species level identification, sequencing of additional loci was performed on representative isolates of the main four taxa of interest in the context of walnut dieback, that is *Neofusicoccum* (n = 6), *Botryosphaeria* (n = 6), *Diaporthe* (n = 6), *Colletotrichum* (n = 6) and *Fusarium* (n = 5; **S1 Table**). For each taxa of interest, phylogenetic reconstructions were performed using concatenated sequence datasets of two to five genes (ITS, translation elongation factor 1-ɑ (EF1-ɑ), β-tubulin (TUB), histone H3 (HIS3), actin (ACT), glyceraldehyde-3-phosphate dehydrogenase (GAPDH), chitin synthase 1 (CHS-1), RNA polymerase largest subunit (RPB1) and RNA polymerase second largest subunit (RPB2)) depending on taxa. The list of genes and reference sequences is provided in **S1 Table**. Overall, the combined sequences of 12 representative Botryosphaeriaceae isolates clustered in two well-supported clades: (i) six isolates clustered together with reference GenBank sequences of *N. parvum* (bootstrap support [BS; %]/Bayesian posterior probability [PP]: 97/1.00), and (ii) six isolates clustered together with reference GenBank sequences of *B. dothidea* (BS/PP: 100/1.00; **S1 Fig**). The combined sequences of 12 representatives *Colletotrichum* isolates also clustered in two well-supported clades: (i) six isolates clustered together with reference GenBank sequences of *C. fioriniae* (BS/PP: 96/1.00), and (ii) six isolates clustered together with reference GenBank sequences of *C. godetiae* (BS/PP: 100/1.00; **S2 Fig**). Last, the combined sequences of the six representative *Diaporthe* isolates and the five representative *Fusarium* isolates all clustered together with reference GenBank sequences of *D. eres* and of *F. juglandicola* in well-supported clades (*Diaporthe*: BS/PP: 99/1.00; **S3 Fig**; and *Fusarium*: BS/PP: 79/0.72; **S4 Fig**).

Each genus was mainly represented by up to two different species in both symptomatic and asymptomatic tissues. *Colletotrichum godetiae* was the major species of the *Colletotrichum* genus (242 out of the 295 collected isolates), as well as *D. eres* belonging to the *Diaporthe* genus (351 out of the 510 collected isolates, the remaining being presumptively *D. foeniculina* based on ITS sequencing although further multilocus sequencing would be needed), and *F. juglandicola* belonging to the *Fusarium* genus (249 out of the 412 collected isolates). Interestingly, *B. dothidea* and *N. parvum* were the only species belonging to the Botryosphaeriaceae family. As the ITS region was insufficient to assign the major part of the isolates belonging to the Didymellaceae family at the genus level, only the *Boeremia* and *Epicoccum* genera were identified, representing 38.1% and 18% of the isolates, respectively.

### Pathogenicity tests

#### Single inoculations on detached twigs

After inoculation on 1-year-old detached twigs of the walnut variety Franquette, all species were able to induce significantly longer necrotic lesions compared to controls (**Fig 3A**). Significant differences in aggressiveness were observed between groups of species (*p* < 0.05), with the mean lesion rate ranging from 0.2 to 13.3 mm/day, with the slowest lesion growth observed for *C. godetiae* (1.0 ± 1.0 mm/day)*, F. juglandicola* (1.1 ± 1.3 mm/day) and *C. fioriniae* (1.3 ± 0.9 mm/day). Botryosphaeriaceae isolates caused the quickest lesion growth (10.1 ± 4 mm/day for *B. dothidea* and 11.1 ± 2.7 mm/day for *N. parvum*), followed by *D. eres*, responsible for moderate lesion growth rate (6.4 ± 2.2 mm/day). Botryosphaeriaceae species were more aggressive and consistently reisolated from inoculated twigs (100% for all isolates; **S2 Table**), followed by *D. eres* (from 44.4% to 100%), while frequency of reisolation ranged from 11.1% to 88.9% for the three other species, questioning their pathogenicity. Positive reisolations were obtained from 16.1% of DJ (drilled but not inoculated twigs covered with petroleum jelly) and DPJ (drilled twigs inoculated with a pure PDA plug and covered with petroleum jelly) control twigs and from 6.7% of D (drilled but not inoculated twigs) control twigs, mainly associated with *Neofusicoccum*, *Diaporthe* and *Alternaria* genera, in line with their presence in asymptomatic samples as highlighting by culture-dependent analyses.

**Fig 3.**
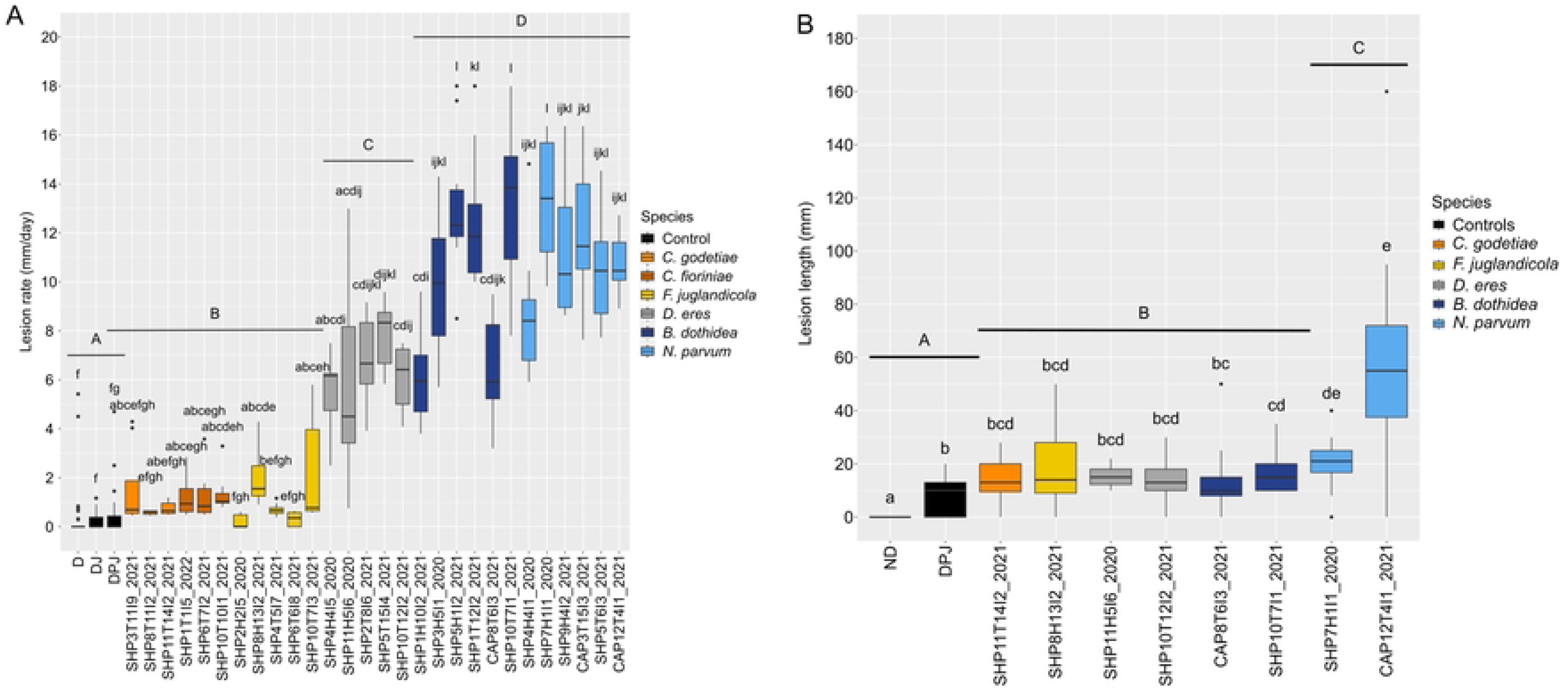
Disease severity of isolates of the six species of interest on 1-year-old English walnut variety Franquette twigs. Mycelium plugs of representative fungal isolates of *Colletotrichum godetiae*, *C. fioriniae*, *Fusarium juglandicola*, *Diaporthe eres*, *Botryosphaeria dothidea* and *Neofusicoccum parvum* were inoculated on (A) detached twigs (lesion rate (mm/day), laboratory conditions) between 10 and 31 days, and (B) attached twigs (lesion length (mm), field conditions) during 6 months. Boxes with the same letter do not differ significantly according to Dunn’s test (ɑ = 0.05). Uppercase letters correspond to comparisons at the species level regardless of isolates. Lowercase letters correspond to comparisons between isolates within a species and the controls. ND: not drilled twigs; D: drilled but not inoculated twigs; DJ: drilled but not inoculated twigs covered with petroleum jelly; DPJ: drilled twigs inoculated with a pure PDA plug and covered with petroleum jelly. All twigs were wrapped with Parafilm.

#### Single inoculations on attached twigs under field conditions

Six months after inoculation on walnut twigs under field conditions, only *N. parvum* isolates and *B. dothidea* isolate SHP10T7I1_2021 caused lesions significantly longer than both ND and DPJ control conditions (*p* < 0.05; **Fig 3B**). The longest lesion length (40.2 ± 35 mm) was caused by *N. parvum* isolates and the species was more consistently reisolated (50% for both isolates, **S2 Table**) compared to *B. dothidea* (only up to 23%), *F. juglandicola* (23%) and *D. eres* (up to 15%). Interestingly, *D. eres* isolate SHP11H5I6_2020 and *C. godetiae* were not reisolated from any of the inoculated twigs, invalidating Koch’s postulates. Reisolation rates of species of interest were obtained from 61.5% of DPJ control twigs, and from 30.8% of ND (not drilled twigs) control twigs although lesion lengths in these samples measured less than 10 mm in average. The same genera as highlighted previously were reisolated from these control conditions.

#### Single inoculation on detached green fruits

The size of lesions observed 12 days after inoculation on green fruits significantly varied depending on the isolate. Only *F. juglandicola* was not able to induce necrosis significantly different from control conditions according to RAUDPC (**Fig 4**). Significant differences in RAUDPC were observed within species (*p* < 0.05). *Neofusicoccum parvum* (RAUDPC = 11.4% and 8.7% for isolates CAP3T15I3_2021 and SHP4H4I1_2020, respectively) and *C. godetiae* (RAUDPC = 7.6%) were the most aggressive, followed by *B. dothidea and D. eres* (RAUDPC = 5.1% and 1.0% for isolates SHP5T15I4_2021 and SHP4H4I5_2020, respectively). Interestingly, intraspecific differences in aggressiveness were observed among *B. dothidea* isolates, with isolate SHP5H1I2_2021 (RAUDPC = 12.2%) being significantly more aggressive than isolate SHP1H10I2_2021 (RAUDPC = 4.4%, *p* < 0.01). Most of the species were reisolated from the margin of necrotic lesions, with reisolation rates ranging from 43% to 100% (**S2 Table**).

**Fig 4.**
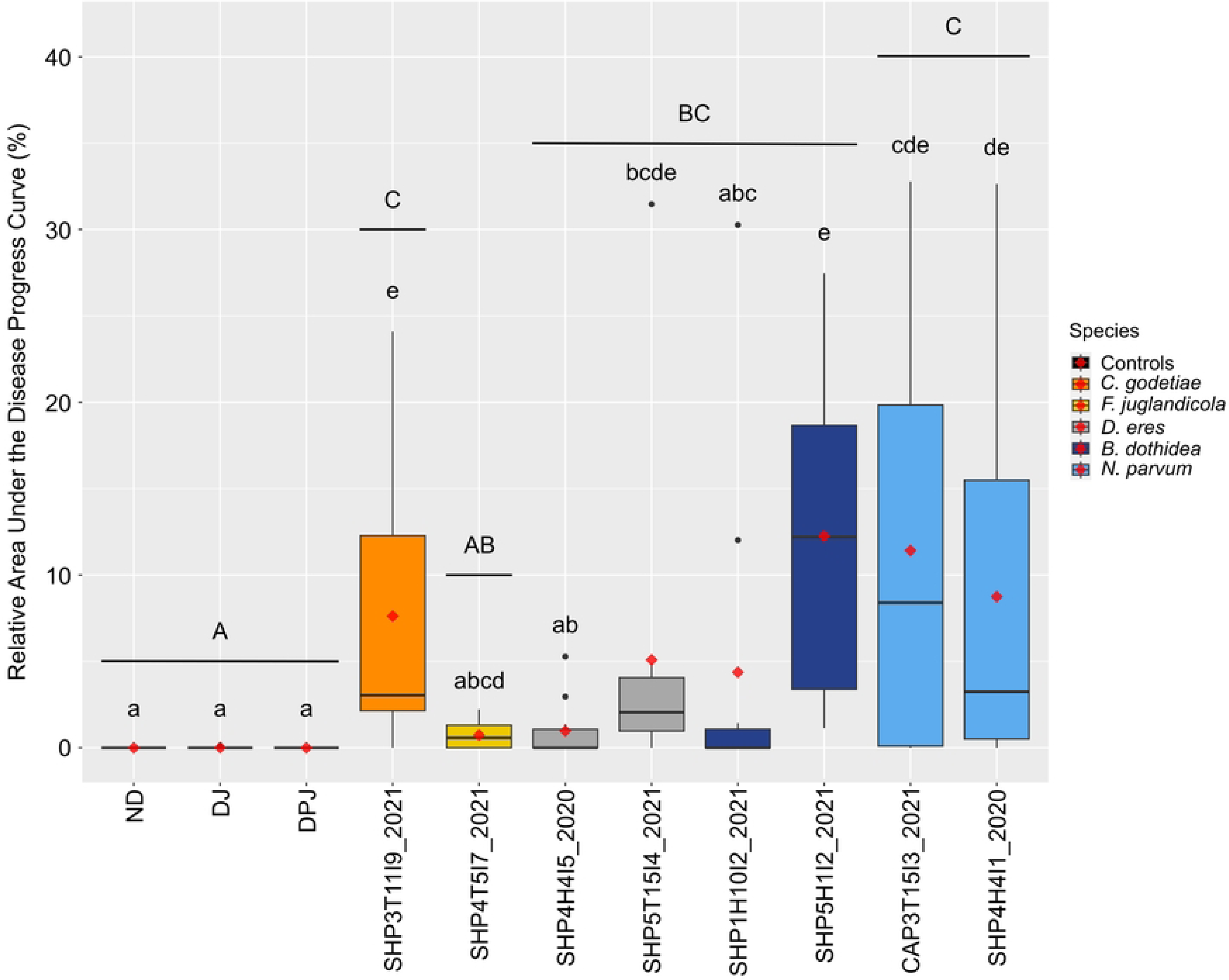
Relative area under the disease progress curve (RAUDPC; %) on detached green fruits of English walnut variety Fernor inoculated with isolates of the six species of interest. Analyses were performed after 12 days after inoculations with mycelium plugs of representative fungal isolates of *Colletotrichum godetiae*, *Fusarium juglandicola*, *Diaporthe eres*, *Botryosphaeria dothidea* and *Neofusicoccum parvum*. Boxes with the same letter do not differ significantly according to Dunn’s test (ɑ = 0.05). Uppercase letters correspond to comparisons at the species level regardless of isolates. Lowercase letters correspond to comparisons between isolates within a species and the controls. ND: not drilled green fruits; DJ: drilled but not inoculated green fruits covered with petroleum jelly; DPJ: drilled green fruits inoculated with a pure PDA plug and covered with petroleum jelly. Red diamonds represent the mean RAUDPC for each isolate.

### Factors impacting the prevalence of pathogenic species

Based on cultural analyses, the prevalence in husks and twigs as well as in symptomatic and asymptomatic samples within each orchard was determined for the six main endophytic/pathogenic species previously described, i.e. *B. dothidea, C. fioriniae, C. godetiae, D. eres, F. juglandicola* and *N. parvum*. The prevalence of these species significantly varied depending on the tissue type and the campaign (**Fig 5**). *Colletotrichum godetiae* and *B. dothidea* were significantly more prevalent in symptomatic walnut husks (ranging from 40% to 70% and 35%, respectively, depending on the year) than in twigs (around 30% and ranging from 17% to 33% depending on the year; *p* = 1.95 × 10^-10^ and *p* = 0.02, respectively). Conversely, *D. eres* and *F. juglandicola* were significantly more prevalent in symptomatic twigs (ranging from 67% to 76% and from 53% to 68% respectively, depending on the year) than in husks (ranging from 44% to 53% and from 25% to 28%; *p* = 1.81 × 10^-7^ and *p* = 4.22 × 10^-13^, respectively). *Neofusicoccum parvum* was similarly prevalent in both types of tissue (with an average of 26% in twigs and 33% in husks, *p* = 0.1) while *C. fioriniae* was poorly prevalent (6% in twigs and 9% in husks, in average, *p* = 0.3). Interestingly, when considering symptomatic and asymptomatic samples, both *F. juglandicola* and *N. parvum* were significantly more prevalent in symptomatic tissues whatever the tissue type (**Fig 6**). The same observation was made for *C. godetiae* but only in walnut husks while the prevalence did not differ significantly between symptomatic and asymptomatic tissues for the three other species (*p* > 0.05, **Fig 6**).

**Fig 5.**
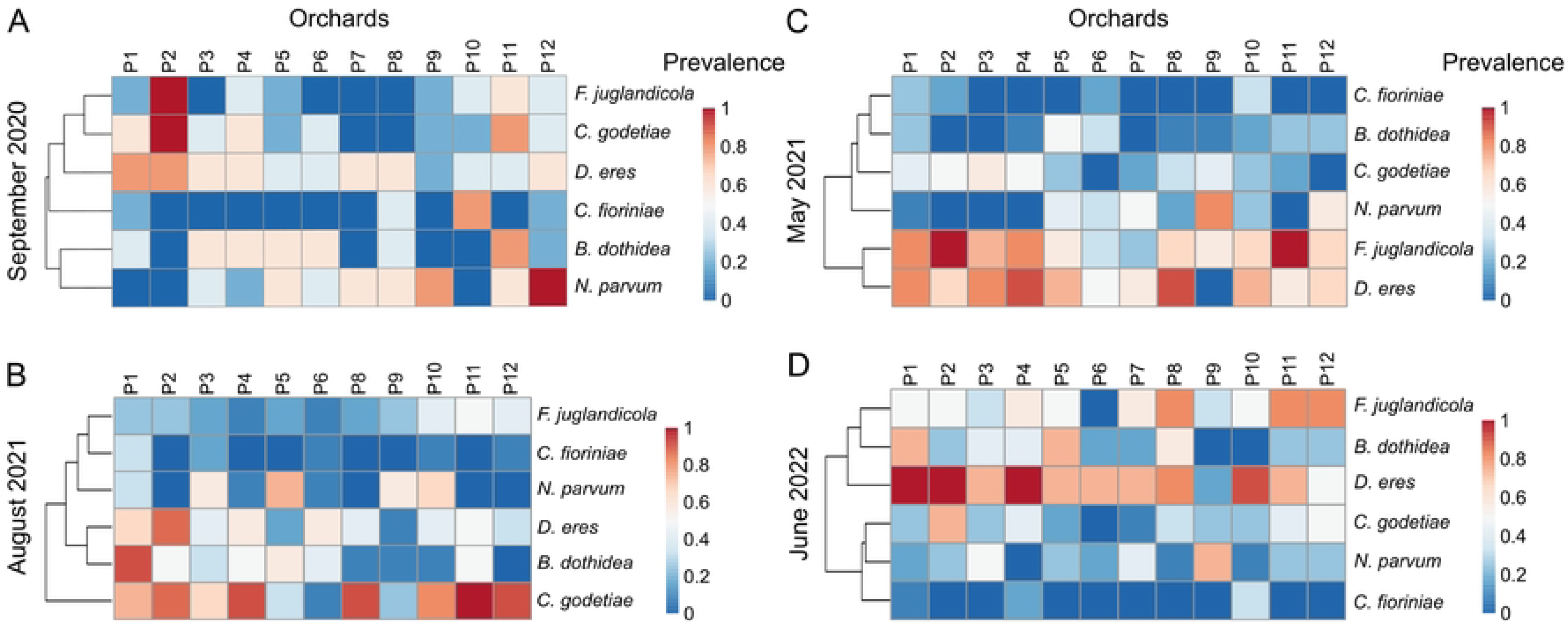
Heatmaps of the prevalence of the six main species of interest on symptomatic tissues. Species of interest were collected from symptomatic walnut husks during (A) September 2020 and (B) August 2021, and from twigs during (C) May 2021 and (D) June 2022.

**Fig 6.**
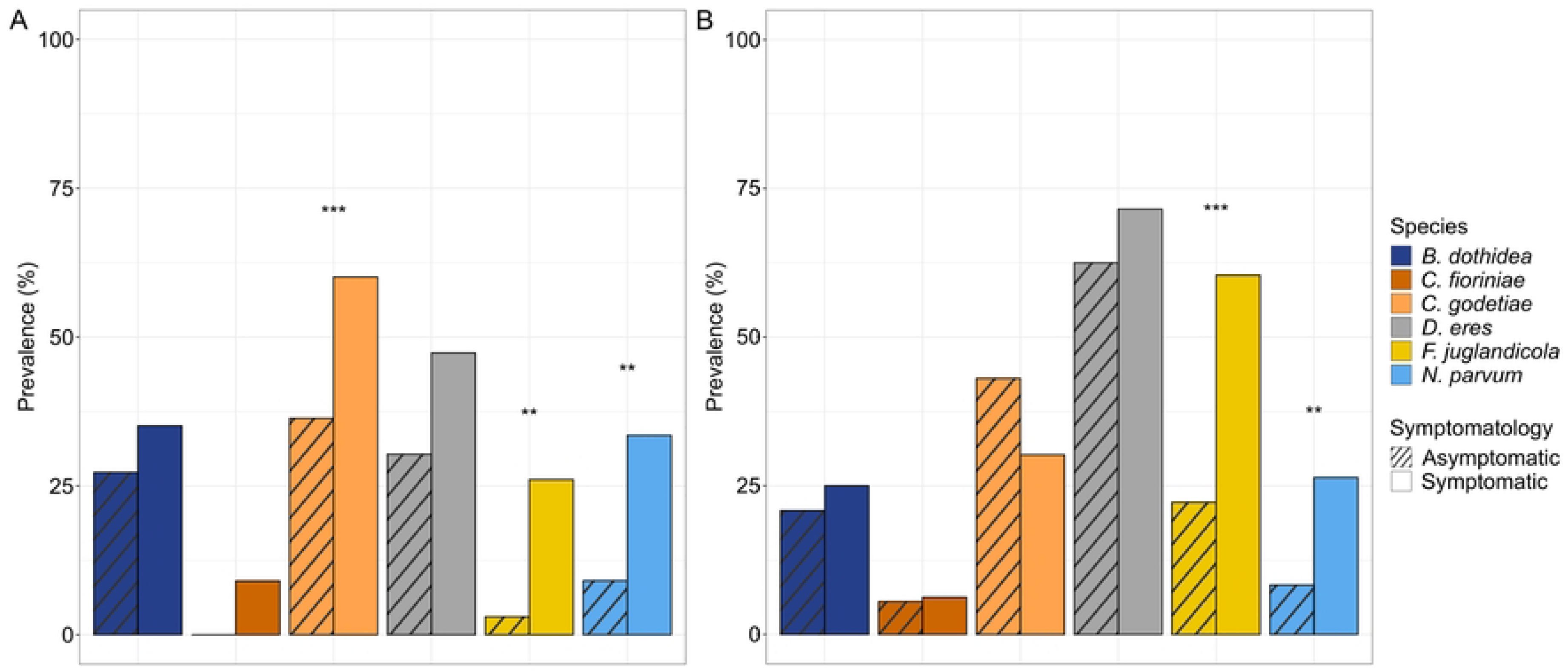
Percentage of asymptomatic and symptomatic tissues contaminated by the six main species of interest out of the total number of tissues analyzed. (A) Walnut husks, (B) walnut twigs. Statistical comparisons were performed per species and per tissue type. “**” and “***” represent highly significant and very highly significant differences based on chi-squared test (ɑ = 0.05).

The PERMANOVA, based on Bray-Curtis distance matrices of prevalence profiles of the 6 species, showed a significant effect of, primarily, the tissue type (husk versus twig; R^2^ = 14.7 %, *p* = 1.0 × 10^-4^), followed by the sampling area (South-East versus South-West; R² = 8.8 %, *p* = 2.3 × 10^-2^). Conversely, the year of sampling did not significantly impact prevalence profiles (*p* > 0.05).

### Description of the fungal pathobiome by ITS2 metabarcoding sequencing

A total of 11,706,886 raw reads of ITS2 were obtained for 282 samples (12 orchards × 3 campaigns × 2 symptomatologies × 3 replicates, and 11 orchards × 1 campaign × 2 symptomatologies × 3 replicates). After read filtering, a large proportion of samples associated with asymptomatic tissues were removed from the dataset due to the absence of fungal reads and a high number of plant reads (i.e. 48 asymptomatic samples preserved out of 141 samples). After this step, a total of 3,887,112 high-quality ITS2 fungal sequences were obtained and clustered into 656 AVSs from 187 samples, including replicates.

The fungal communities in both symptomatic walnut husks and twigs were predominantly composed of Ascomycota (99.1% and 99.7%, respectively) followed by Basidiomycota. Although the taxonomic resolution of ITS2 does not allow differentiating between *Botryosphaeria* and *Neoscytalidium* [35], results of the culture-dependent analysis with the unique isolation of *B. dothidea*, strongly suggest that ASVs assigned to *Botryosphaeria/Neoscytalidium* very likely belong to the *Botryosphaeria* genus. The genera most frequently encountered by culture-dependent approach (*Botryosphaeria, Colletotrichum, Diaporthe, Fusarium* and *Neofusicoccum*) were also found to be predominant in symptomatic samples through metabarcoding analysis (**Fig 7**). These genera represented between 20.9% and 99.3% of total reads in symptomatic walnut husks and twigs in an orchard- and campaign-dependent manner (**Fig 7**). The genus *Plectosphaerella* was also among the top 15 fungal genera in walnut husks even if associated with few orchards (four orchards collected in 2020 and two collected in 2021, respectively), as well as *Gibellulopsis* which was represented in the majority of orchards collected in both years by less than 1% to 75% relative abundance, and *Vishniacozyma*, represented by relatively low relative abundance in all the orchards collected in 2020 (from less than 1% to 1.7% of relative abundance) and in only four orchards collected in 2021 (P1, P2, P10 and P12). In symptomatic twigs, the *Boeremia* and *Angustimassarina* genera were also abundant (from less than 1% to 57.3%, and less than 1% to 25.3%, respectively) in the majority of orchards across both years. These genera, as well as the less abundant genera (e.g. *Cladosporium*, *Ophiognomonia* and *Juglanconis*) have never been collected with the culture-dependent method, or only to a very small extent.

**Fig 7.**
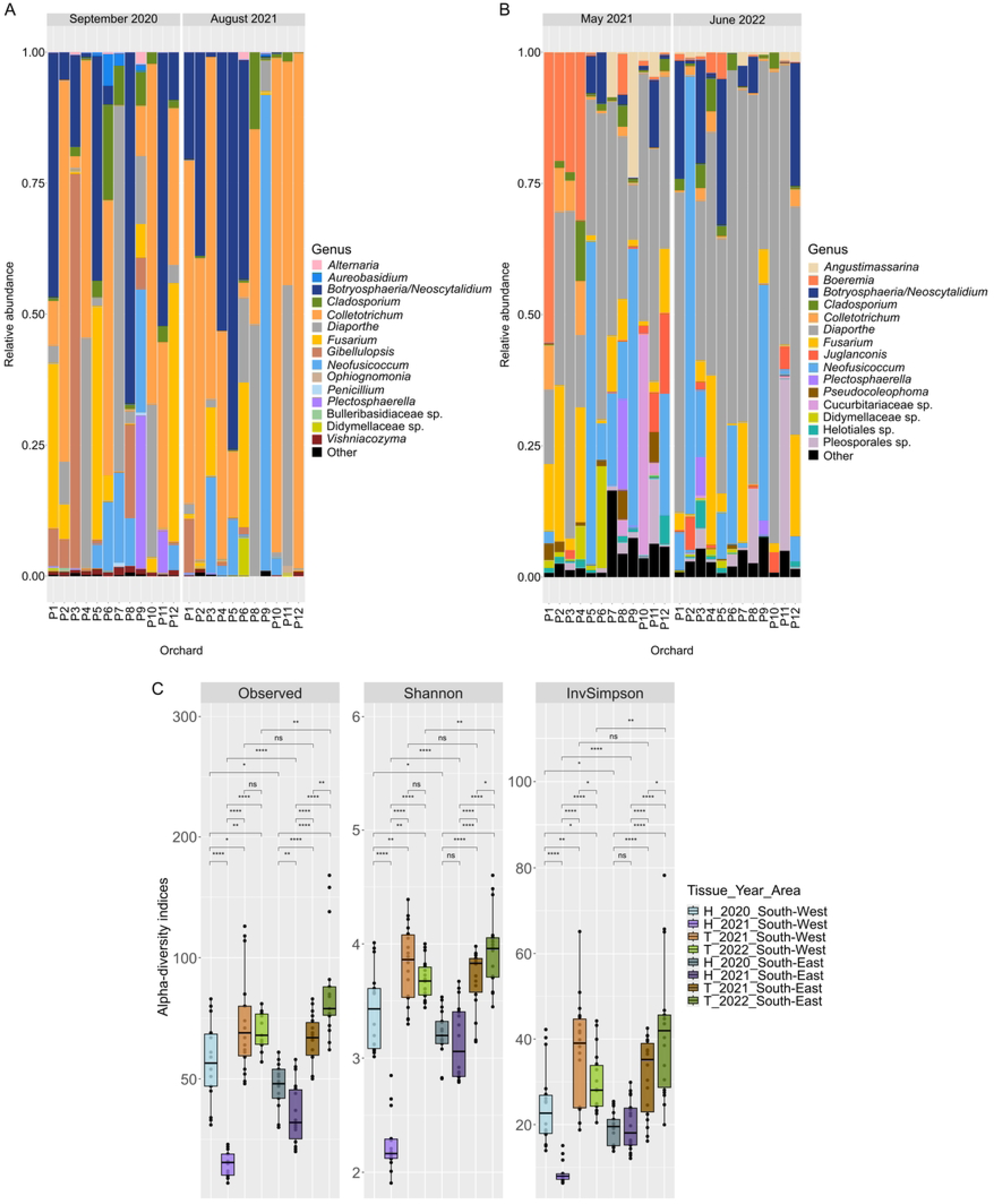
Relative abundance of the 15 predominant genera from symptomatic tissues and α-diversity indices based on ITS2 metabarcoding. (A) Walnut husks, (B) walnut twigs. (C) Αlpha-diversity indices of the combination of symptomatic tissue type (H: Husks, T: Twigs), sampling year and area calculated based on the rarefied dataset. “ns”, “*”, “**” and “***” represent non-significant, significant, highly significant and very highly significant differences based on independent t.test (ɑ = 0.05).

Interestingly, when comparing relative abundances in symptomatic husks versus twigs, some genera were significantly more abundant in husks: *Aureobasidium* (log2 fold change |3|, *p* < 0.01)*, Botryosphaeria* (log2 fold change |3|, *p* < 0.001)*, Colletotrichum* (log2 fold change |6|, *p* < 0.001)*, Gibellulopsis* (log2 fold change |6|, *p* < 0.001) and *Vishniacozyma* (log2 fold change |1|, *p* < 0.05). Conversely, the relative abundance was significantly higher in symptomatic twigs than in husks for *Juglanconis* (log2 fold change |10|, *p* < 0.001), *Diaporthe* (log2 fold change |1|, *p* < 0.001)*, Boeremia* (log2 fold change |7|, *p* < 0.001) and *Angustimassarina* (log2 fold change |10|, *p* < 0.001). Noteworthy, the *Colletotrichum* genus also appeared to be significantly more abundant in symptomatic walnut husks collected in August 2021 than in those collected in September 2020 (log2 fold change |2|, *p* < 0.001). The analysis of the few asymptomatic samples also led to the detection of *Colletotrichum*, *Diaporthe* and *Ophiognomonia* and *Vishniacozyma* genera in husks and *Angustimassarina, Boeremia, Botryosphaeria, Diaporthe, Fusarium* and *Neofusicoccum* in twigs.

Alpha-diversity indices, that is, observed richness (number of observed ASVs) and the Shannon and Simpson’s inverse (InvSimpson) indices, were significantly higher in symptomatic twigs compared to symptomatic husks, regardless of sampling campaign or areas (**Fig 7C**). These indices were also significantly different according to sampling campaigns and areas. In particular, walnut husks collected in the South-West in September 2020 showed significantly higher richness, diversity and equitability than those collected in the same region in August 2021 while twigs collected in June 2022 showed significantly higher indices values than those collected in May 2021. Overall, walnut husks or twigs were significantly more diverse and richer in the South East than their counterparts collected in the South West, except for husks collected in September 2020 (**Fig 7C**).

The PERMANOVA, based on Bray-Curtis distances, showed that fungal communities in symptomatic samples were first impacted by tissue types (R² = 16.6 %, *p* = 1.0 × 10^-4^), followed by sampling year (R^2^ = 11,6 %, *p* = 1.0 × 10^-3^) and sampling area (R² = 4.6 %, *p* = 2.0 × 10^-4^).

### Effect of climatic factors on fungal pathobiome composition and structure

To better understand the impact of meteorological factors promoting the major identified genera, daily temperatures and rainfall were retrieved from 6 weather stations, evenly located in the South-East and in the South-West (**S3 Table**). The data were collected from April to August in 2020, 2021 and 2022, that is from early springs to late summers. Since mean temperature or rainfall did not differ significantly between the three weather stations within a production area, regardless of the month and year, data from the three weather stations were combined. Overall, whatever the production area, April, May, July and/or August were associated with significantly lower mean temperatures in 2021 than in 2020 or 2022, only June was significantly hotter in 2021 than in 2020. In addition, mean accumulated rainfall was significantly higher in May and in July 2021 than in 2020 or 2022, regardless of the production area (**S5 and S6 Figs** and **S4 Table**). At the geographic level, mean temperature and rainfall were generally higher in the South-West than in the South-East, with significant differences in terms of mean temperature in April 2021, May 2020 and May 2022, and mean rainfall in April 2020 (**S4 Table**)

To go further and assess potential correlations between meteorological variables and the prevalence of the six main species, a canonical correspondence analysis (CCA) was performed (**Fig 8**). As can be seen from the length of the arrows, mean monthly temperature and cumulative monthly rainfall both had a strong influence on the prevalence of fungal species (**Fig 8**). Moreover, the orientation of the arrows indicated that the two variables were negatively correlated, as confirmed by Pearson’s correlation test (corr = -0.81, *p* =3.1 × 10^-12^). Among species of interest, *N. parvum* was positively influenced by temperature and negatively influenced by cumulative rainfall, indicating a higher prevalence of this species associated with warm and dry weather. The same was also observed for *D. eres*, although to a lesser extent. Conversely, the two species belonging to the *Colletotrichum* genus were positively influenced by cumulative rainfall, and negatively influenced by temperature, indicating that these species were more prevalent during sampling campaigns associated with wet and cold weather. The two other species showed opposite correlations. While *B. dothidea* was positively influenced by cumulative rainfall and to a lesser extent by temperature, *F. juglandicola* was negatively influenced by cumulative rainfall, and its prevalence did not seem to be influenced by the temperature.

**Fig 8.**
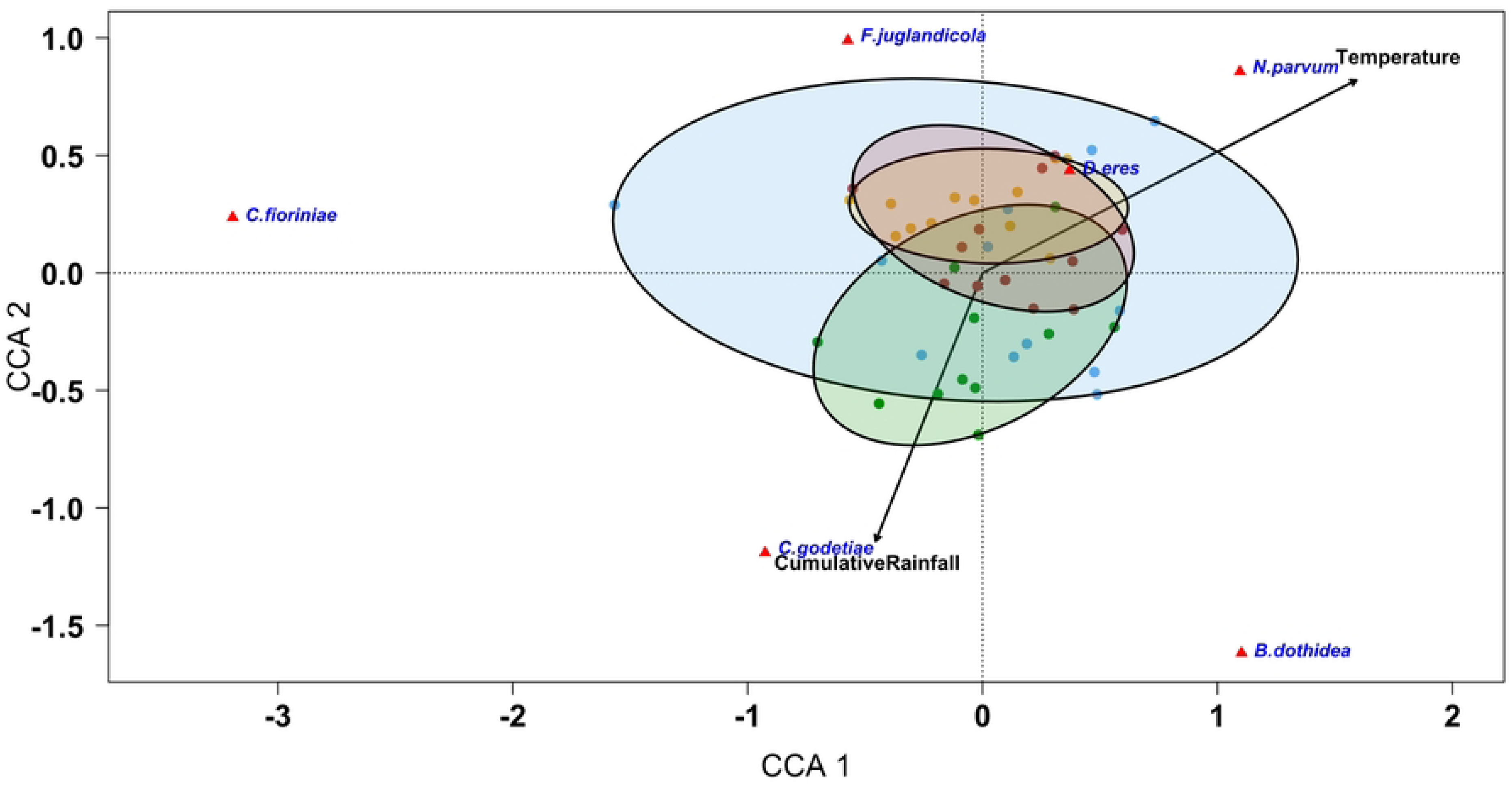
Ordination diagram of canonical correspondence analysis based on the prevalence of the six main fungal species associated with walnut dieback symptoms with two explanatory weather variables over the three years of sampling (from 2020 to 2022). Red triangles represent fungal species and black arrows represent weather variables (Cumulative rainfall (mm) and temperature (°C)). Colored points represent the orchards sampled during the four sampling campaigns: symptomatic walnut husks collected in September 2020 (blue) and in August 2021 (green), and symptomatic twigs collected in May 2021 (yellow) and in June 2022 (brown).

### Analysis of the co-occurrence and association patterns within the phytomicrobiome

#### Culture-dependent based analyses

First, pairwise Pearson’s correlations were calculated based on the prevalence of the six main species isolated from symptomatic husks and twigs at the orchard level. In both tissues, *N. parvum* and *D. eres* were significantly negatively correlated (Pearson’s correlation coefficient (PCC) = -0.71 and PCC = -0.91, respectively; **Fig 9A,B**) while *C. godetiae* and *F. juglandicola* were significantly positively correlated (PCC = 0.73 and PCC = 0.63, respectively; **Fig 9A,B**). Finally, a significant negative correlation was evidenced between *N. parvum* and *C. godetiae* in symptomatic walnut husks (PCC = -0.68; **Fig 9A**).

**Fig 9.**
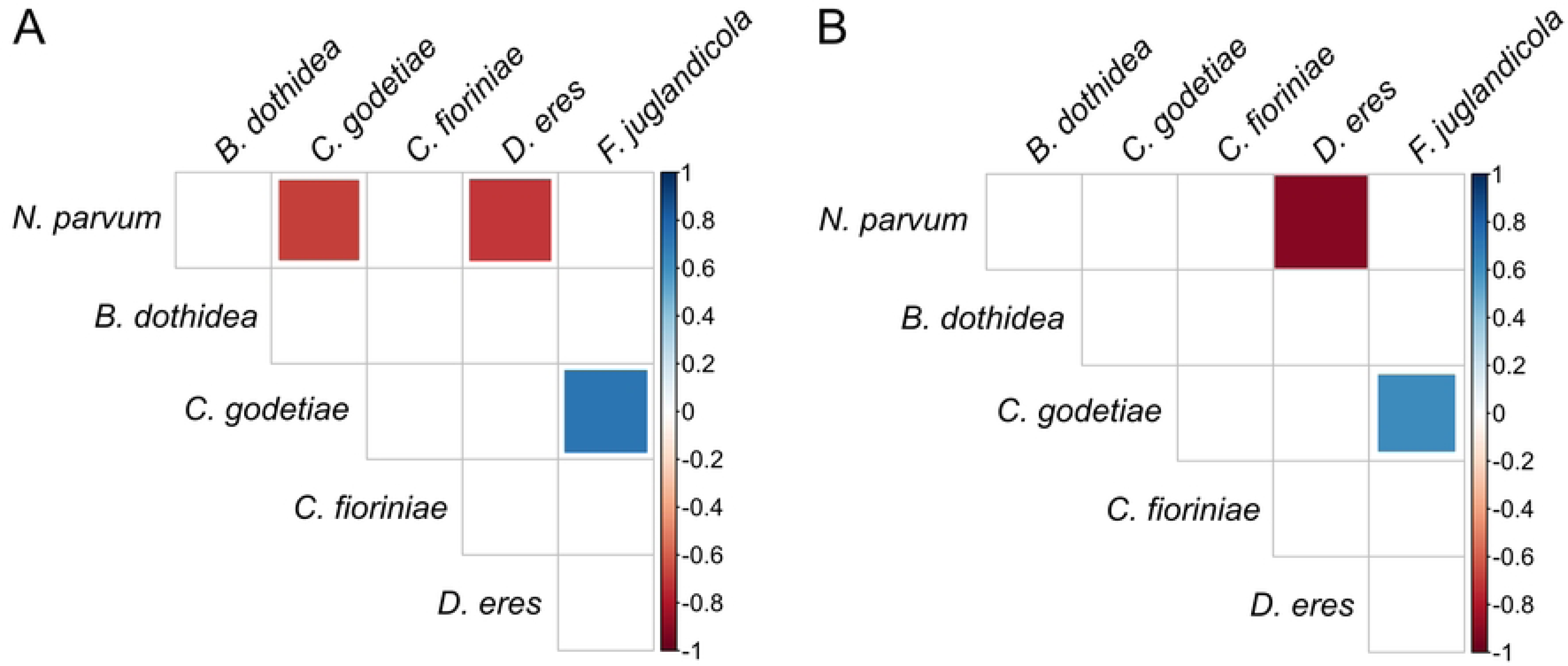
Correlogram representing Pearson’s correlation coefficient between the 6 main species of interest based on their prevalence on symptomatic tissues. (A) Walnut husks, (B) walnut twigs. Only significant correlations (*p* < 0.05) are plotted.

Second, chi-squared tests were computed to determine whether a species was more frequently isolated alone or in association at the tissue level. The frequencies of dual association at the tissue level were calculated considering the number of tissues contaminated by either or both species regardless of the other species. Interestingly, all the species of interest were more frequently found in association with at least one species than alone (from 2 to 16 times more in symptomatic tissues and from 1 to 14.5 times in asymptomatic ones). In symptomatic tissues, among all species, *N. parvum* was the species most frequently isolated alone either in husks or twigs (33.3% of husks and 30.3% of twigs contaminated with *N. parvum* were exclusively contaminated with this species), followed by *D. eres* in twigs (17.4%) and *C. godetiae* in husks (12.5%). Interestingly, *B. dothidea* and *F. juglandicola* were found alone in only 1.4% and 6.4% of symptomatic twigs associated with these species, respectively. In both tissues, the two Botryosphaeriaceae species were not frequently simultaneously isolated. This co-occurrence represented 17.3% of husks and 10.4% of twigs contaminated by these species, while *N. parvum* was more frequently isolated without *B. dothidea* (81.6% twigs and 69.8% of husks) and vice versa (i.e. *B. dothidea* without *N. parvum* found in 80.5% twigs and 71% of husks). Furthermore, the most frequent dual association was that of *C. godetiae* and *D. eres* in symptomatic husks (12.9% of husks were contaminated with *C. godetiae* and *D. eres* exclusively), and that of *F. juglandicola* and *D. eres* in symptomatic twigs (13.8% of twigs were contaminated with *F. juglandicola* and *D. eres* exclusively). Moreover, the latter dual association was significantly more detected in all symptomatic twigs contaminated by *D. eres* and/or *F. juglandicola*, whatever the other species, than in husks under the same conditions (51% vs 21%, *p* = 2.5 × 10^-6^), and, in symptomatic twigs, the two species were significantly more associated with each other regardless of the presence of other species (51%) than without each other (31% with *D. eres* and without *F. juglandicola* whatever the other species, *p* = 2.1 × 10^-4^; and 18% with *F. juglandicola* and without *D. eres* whatever the other species, *p* = 2.1 × 10^-13^). Interestingly, the tripartite association of *D. eres, C. godetiae* and *F. juglandicola* without other species was the most frequently encountered and represented 8.2% of symptomatic twigs contaminated by these species.

When confronting the results based on comparisons of prevalence to those based on associations in symptomatic tissues, the positive correlation between *C. godetiae* and *F. juglandicola* based on prevalence data was also confirmed at the tissue level for twigs only. The two species were more frequently found associated with each other (34.7% of total twigs contaminated by *C. godetiae* and/or *F. juglandicola*) than *C. godetiae* without *F. juglandicola* (10.4%), regardless of the presence of the other species (*p* = 3.1 × 10^-7^). Likewise, the negative correlations between *N. parvum* and *D. eres* in twigs and husks was confirmed at the tissue level for husks only since *D. eres* and *N. parvum* were significantly more frequently isolated separately (in 53.7% and 34.6% of husks contaminated with at least one of the two species, respectively) than associated with each other (11.7%, *p* = 6.8 × 10^-12^ and *p* = 2.4 × 10^-4^, respectively). Finally, *C. godetiae* was significantly more frequently isolated without *N. parvum* (in 58% of husks contaminated with at least one of the two species) than in association with *N. parvum* (16.7%) whatever the other species (*p* = 9.9 × 10^-13^), confirming the negative correlation between *C. godetiae* and *N. parvum* in symptomatic husks **(Fig 9A)**.

#### Metabarcoding-based co-occurrence analyses

In total, six positive and seven negative significant correlations were evidenced between nine genera out of the 15 top genera associated with symptomatic walnut husks (**Fig 10A**). No positive or negative significant correlations were found between the main pathogenic genera of interest (i.e. *Colletotrichum*, *Diaporthe* and *Neofusicoccum*). Nevertheless, a positive significant correlation was highlighted between *Neofusicoccum* and *Aureobasidium* (corr = 0.32), as well as a negative significant correlation between *Colletotrichum* and *Aureobasidium* (corr = -0.31; **Fig 10A**).

**Fig 10.**
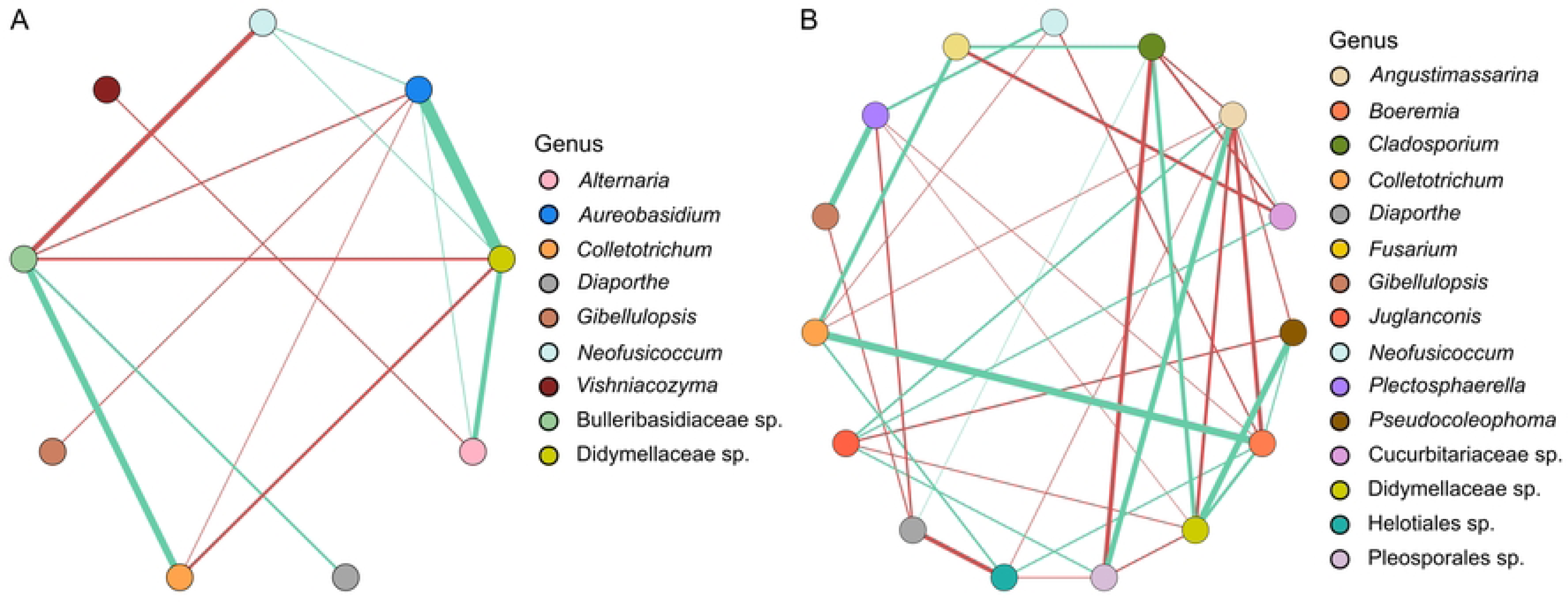
SparCC correlation networks observed between the main taxa associated with symptomatic tissues based on metabarcoding sequencing. (A) Walnut husks, (B) walnut twigs. The relative abundance of each taxon was calculated by summing the abundance of each corresponding ASV. Nodes correspond to taxa, and connecting edges indicate correlation between them. Only nodes associated with a correlation strength value greater than 0.3 (positive correlation, green edges) and less than -0.3 (negative correlation, red edges) were plotted. Only significant correlations (*p* < 0.05) were plotted.

In twigs, 17 out of 37 correlations were significantly positive (**Fig 10B**). Interestingly, a positive significant correlation was detected between *Colletotrichum* and *Fusarium* (corr = 0.52) as observed based on culture-dependent analyses, and between *Colletotrichum* and *Boeremia* (corr = 0.70), while *Colletotrichum* and *Neofusicoccum* were negatively correlated (corr = -0.32; **Fig 10B**), also in line with our previous result.

## Discussion

A total of 12 walnut orchards from the two main production areas in France, selected on the basis of frequent observation of dieback symptoms on English walnut trees, were surveyed during four sampling campaigns. The observed symptoms included defoliation of twigs, dead buds and internal wood discoloration, as well as fruit necrosis, blight and mummification. This research represents the first comprehensive study providing the etiology and analysis of fungal communities and their interactions in walnut trees displaying these symptoms in France, using both cultural and metabarcoding approaches.

The major part of the isolates recovered from walnut husks and twigs, either symptomatic or asymptomatic, were identified based on their morphological characteristics and sequencing. The following species were identified: *Botryosphaeria dothidea* and *Neofusicoccum parvum* within the Botryosphaeriaceae family, *Diaporthe eres* within the *Diaporthe* genus, *Colletotrichum fioriniae* and *C. godetiae* within the *Colletotrichum* genus, and *Fusarium juglandicola* within the *Fusarium* genus.

*Diaporthe eres* and *F. juglandicola* were consistently the most frequently isolated fungi from symptomatic tissues, followed by *Colletotrichum* isolates (mainly represented by *C. godetiae* and *C. fioriniae* to a lesser extent) and isolates from the Botryosphaeriaceae family, exclusively represented by *B. dothidea* and *N. parvum*. These results were confirmed by metabarcoding, revealing the presence of the same genera in both symptomatic walnut husks and twigs. This range of diversity associated with the disease was also evidenced in Californian walnut orchards [36]. However, only Botryosphaeriaceae and/or *Diaporthe* species were reported in similar symptomatic walnut tissues in other studies conducted in Australia [22], California [8,20], Chile [23], China [24], Italy [11] and Hungary [13]. Interestingly, a higher level of diversity was found in Spain where *Cytospora* species were also isolated in addition to these two fungal groups [10], as well as Diatrypaceae, Togniniaceae and *Cadophora* species in the Czech Republic [9]. Moreover, most of these studies indicate that Botryosphaeriaceae isolates were more frequently collected than *Diaporthe* ones [8–10,22,23], unlike what we found during the four collection campaigns in France. Surprisingly, in French orchards, only two genera of Botryosphaeriaceae were identified with both culture-dependent and - independent approaches. In contrast, a much higher level of diversity was found in California where the disease was first observed in the early 2000s and where six genera are consistently reported [8], and, to a lesser extent, in Spain (four genera) [10]. A recent study of the populations of the two species in French walnuts revealed contrasting genetic patterns with *N. parvum* populations from Californian and Spanish walnuts but a higher genetic similarity with populations from French grapevines suggesting a potential transfer between hosts at a more local scale [37]. The two Botryosphaeriaceae species as well as *D. eres* have been previously reported to cause walnut branch dieback and fruit necrosis and blight worldwide [8–11,13,38,39]. The three other species reported in French walnut orchards were not commonly associated with walnut dieback in other countries. The *Colletotrichum* species are mainly associated with walnut anthracnose disease on fruits [5], but one recent study has reported twig blight symptoms caused by *C. godetiae* in Hungary [40]. Regarding *F. juglandicola*, no study has currently indicated its involvement in branch dieback and fruit necrosis and blight of walnut trees, probably due its recent description and identification [41] .

Among the six main species associated with the symptoms, Botryosphaeriaceae species were the most aggressive in all tissues evaluated (detached and attached twigs and detached fruits), followed by *D. eres*. Moreover, *C. godetiae* was also part of the most aggressive species but only on detached fruits. No significant difference in aggressiveness was evidenced between *B. dothidea* and *N. parvum* when inoculated on detached twigs under laboratory conditions, as obtained by López-Moral *et al.* (2020) [10]. Nevertheless, *N. parvum* was more aggressive than *B. dothidea* when inoculated on attached twigs under orchard conditions. Numerous studies have already shown that *N. parvum* is one of the most aggressive Botryosphaeriaceae species and *B. dothidea* one of the least aggressive when inoculated *in planta* [8,10,11]. The same conclusions were obtained when these species were inoculated on detached fruits under laboratory conditions [10,11], as shown here on variety Fernor. In agreement with our results, *Diaporthe* species generally cause less severe symptoms than *Neofusicoccum* species [8,10,23]. The study conducted by Varjas *et al.* (2020) [40] reported necrotic lesion length caused by *C. godetiae* between 12 and 17 mm after 14 days of experimentation, i.e. between 0.8 and 1.2 mm per day, in the same order of magnitude as our own observations. Finally, the high aggressivity of *C. godetiae* on detached fruits is consistent with the study of Da Lio *et al.* (2018) [5], and with its ability to cause anthracnose symptoms on these tissues.

All these species were collected from both symptomatic and asymptomatic tissues, highlighting a potential endophytic lifestyle and latent infections, as consistently reported by many studies. The significantly higher prevalence on symptomatic than on asymptomatic walnut husks of *C. godetiae,* and on both organs for *N. parvum* and *F. juglandicola* indicate that these species could play a more or less important role in symptom development, depending on their pathogenicity and the interactions in which they are involved. For example, *Colletotrichum* spp., responsible for walnut anthracnose and more abundantly isolated from husks, could create lesions that would enable other species to penetrate inside the fruit or weaken the tree, facilitating further infections. The aggressive species *N. parvum* is more likely to directly cause symptoms in husks and twigs when present while *F. juglandicola,* associated with the lowest disease severity, if any, likely plays a secondary role in disease development, probably as an opportunistic agent. Concerning *D. eres,* its prevalence of over 60% in both symptomatic and asymptomatic twigs in addition to its moderate to high aggressiveness suggest a participation of this species to symptom development in twigs as well as its potential endophytic lifestyle. The absence of significant difference of prevalence in asymptomatic and symptomatic tissues contaminated by *B. dothidea* associated with a known moderate aggressiveness rather suggest a moderate involvement in disease development.

Our study revealed that the main factor influencing the prevalence profile of the six main species in symptomatic tissues was the tissue type. *Botryosphaeria dothidea* and *C. godetiae* were significantly more associated with husks, while *D. eres* and *F. juglandicola* were more frequently associated with twigs, in agreement with our metabarcoding-based results. The year of sampling also influenced the variation of mycobiota composition, affecting both prevalence and relative abundance. The sampling area also showed a significant impact on the structure of the fungal communities, as well as meteorological conditions, such as temperature and rainfall. Thus, the terroir could also play a role in the composition of fungal communities. Overall, the differences in meteorological conditions between the three years studied highlighted a significantly higher prevalence of *D. eres* and *N. parvum* in samples collected during warmer and dryer years, contrary to *C. fioriniae* and *C. godetiae*. Nevertheless, the seasons studied were either warmer and drier or colder and wetter, making it challenging to distinguish the individual effects of these two parameters (i.e. temperature and rainfall) on the abundance of these organisms. In the literature, it has been shown that infections caused by *Colletotrichum* species generally occur in rainy, humid and warm weather [42–45]. Interestingly, in other studies, the Botryosphaeriaceae and *Diaporthe* species infections have been associated with warm temperatures and abundant rainfall [46,47]. However, the optimal growth temperature is reported to be generally lower for *Diaporthe* and *Colletotrichum* than for Botryosphaeriaceae species [8,10,48,49]. Meteorological conditions are also known to influence spore dissemination of Botryosphaeriaceae, *Diaporthe* and *Colletotrichum* species, primarily through rain-disseminated spores [47,50], as well as to affect the physiological state of plants, notably during hot periods, which could also facilitate the development of pathogenic species and thus plant disease epidemics [51,52].

This study also highlighted potential interactions between the main fungal pathogenic species. First, we found that the prevalence in orchards and presence at the tissue level of *N. parvum* were both negatively correlated to those of *D. eres* in both symptomatic husks and twigs. Interactions between one species of *Neofusicoccum* (*N. mediterraneum*) and one species of *Diaporthe* (*D. rhusicola*, syn. *D. foeniculina*) was previously investigated by Agustí-Brisach *et al.* (2019) [20], and evidenced a strong negative correlation between these two species. Additionally, they demonstrated an inhibitory effect of *N. mediterraneum* spores on the germination of *D. rhusicola* spores. An additional negative correlation was highlighted between *N. parvum* and *C. godetiae* on symptomatic walnut husks, with the latter being significantly more frequently isolated in the absence of *N. parvum* than in its presence. This negative correlation is most likely explained by climatic conditions favorable to *Colletotrichum* in 2021 and to Botryosphaeriaceae in 2020. Finally, the positive correlation between *C. godetiae* and *F. juglandicola* on both symptomatic walnut husks and twigs, based on both prevalence estimations and SparCC networks, prompted us to hypothesize that the presence of one species facilitated the development of the other and possibly, on husks, lesions caused by *Colletotrichum* could be further colonized by *F. juglandicola*. Additional correlations were highlighted between the main pathogenic species and other members of the pathobiome. Among them, a positive correlation between *Neofusicoccum* and *Aureobasidium* genera was detected based on co-occurrence networks on symptomatic walnut husks, as well as a negative correlation between *Colletotrichum* and *Aureobasidium*. The genus *Aureobasidium*, in particular the endophytic species *Aureobasidium pullulans*, has proven useful as a biocontrol agent against many fungal pathogens such as *Botrytis cinerea* [53], *Colletotrichum acutatum* [53] and *Diplodia seriata* [54]. As a consequence, the abundance of this genus could be either promoted by the presence of its target, leading to positive correlation or the control of the target could lead to a negative correlation; these hypotheses should be further tested. A transcriptomic study conducted on liquid co-cultures of *A. pullulans* and *Fusarium oxysporum* highlighted the up-regulation of many genes of *A. pullulans* potentially involved in a biocontrol activity in response to *F. oxysporum* presence according to the authors [55]. Secondly, *Aureobasidium* could also play a role in symptom development, as hypothesized by Nerva *et al.* (2022) [56]. Analysis of transcriptomes in grapevine wood evidenced an up-regulation of genes involved in carbohydrate degradations and in transporters together with a high abundance of *N. parvum* and *A. pullulans,* which could suggest an involvement of these two species in symptom development [56]. In symptomatic walnut twigs, a positive correlation was also detected between *Colletotrichum* and *Boeremia*. This positive correlation involved one of the main genera of our study, and secondary pathogens, identified with the culture-independent method. Indeed, *Boeremia* species (family Didymellaceae), mainly *B. exigua*, was already reported as responsible for branch blight [57] and leaf spot [28,58] on walnuts. Thus, the development and infections caused by some pathogens of walnut trees could facilitate the development of others, as illustrated previously. These results highlight how crucial it is to deepen our knowledge of the interactions between members of the pathobiome, this disease resulting from a complex of fungal pathogenic species.

To our knowledge, this study is the first to explore the fungal pathobiome associated with branch dieback and fruit blight and necrosis of walnut trees using complementary cultural and metabarcoding approaches. The six main species *B. dothidea, C. fioriniae, C. godetiae, D. eres, F. juglandicola* and *N. parvum* were identified as associated with symptomatic tissues, although only the Botryosphaeriaceae and Diaporthaceae species were able to induce symptoms on walnut tissues, while the *Colletotrichum* species were only aggressive on detached fruits. Based on a comprehensive analysis of co-occurrence data at field and tissue levels under field conditions, this study also provides first clues towards a better understanding of the interactions between members of the fungal pathobiome during disease development.

## Materials and methods

### Sampling sites and sample collection

From late summer 2020 to early summer 2022, field surveys were conducted to study the occurrence of branch dieback and fruit blight and necrosis of English walnut in southern France. Twelve commercial orchards (P1 to P12) belonging to the Auvergne-Rhône-Alpes region (South-East production area, P1 to P6, Franquette variety) and the Nouvelle-Aquitaine region (South-West production area, P7 to P12, variety Franquette or cultivar Serr) were surveyed (**Table 1**). In each orchard, symptomatic and asymptomatic walnut tissues were collected from different trees during four sampling campaigns. Walnut husks were collected in September 2020 and August 2021, and twigs were collected in May 2021 and June 2022. In each orchard, five symptomatic walnut husks were collected during September 2020, while 12 symptomatic and three asymptomatic walnut husks or twigs were collected during the three other campaigns, when possible. The P7 orchard could not be sampled during the sampling campaign of August 2021. All samples were stored at 4°C until analyzed within one week after sampling.

**Table 1.**
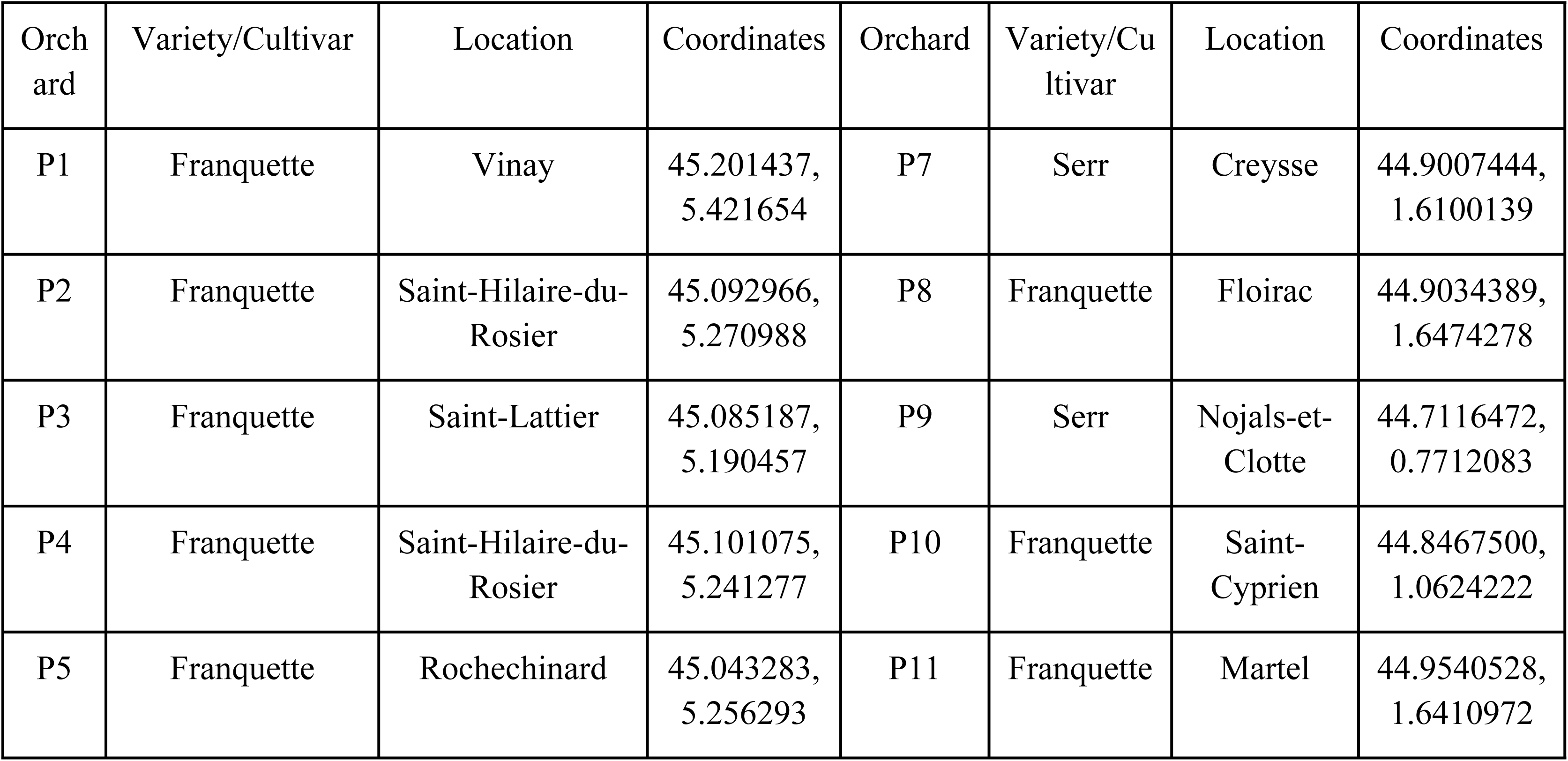
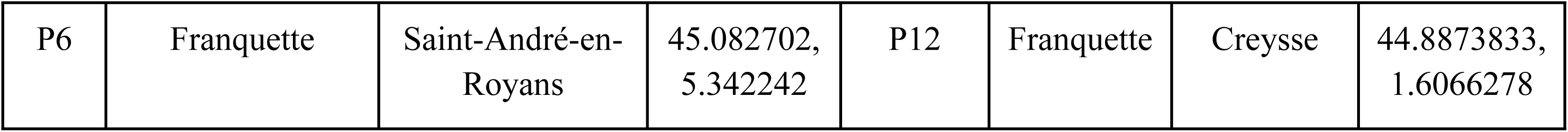
Description of walnut orchards sampled for this study.

| Orchard | Variety/Cultivar | Location | Coordinates | Orchard | Variety/Cultivar | Location | Coordinates |
| --- | --- | --- | --- | --- | --- | --- | --- |
| P1 | Franquette | Vinay | 45.201437,<br>5.421654 | P7 | Serr | Creysse | 44.9007444,<br>1.6100139 |
| P2 | Franquette | Saint-Hilaire-du-Rosier | 45.092966,<br>5.270988 | P8 | Franquette | Floirac | 44.9034389,<br>1.6474278 |
| P3 | Franquette | Saint-Lattier | 45.085187,<br>5.190457 | P9 | Serr | Nojals-et-Clotte | 44.7116472,<br>0.7712083 |
| P4 | Franquette | Saint-Hilaire-du-Rosier | 45.101075,<br>5.241277 | P10 | Franquette | Saint-Cyprien | 44.8467500,<br>1.0624222 |
| P5 | Franquette | Rochechinard | 45.043283,<br>5.256293 | P11 | Franquette | Martel | 44.9540528,<br>1.6410972 |
| P6 | Franquette | Saint-André-en-Royans | 45.082702,<br>5.342242 | P12 | Franquette | Creysse | 44.8873833,<br>1.6066278 |

### Collection of fungal isolates and evaluation of prevalence

Environmental samples were sterilized and cut into small pieces as described in Belair *et al.* (2023) and 5 pieces of each sample were placed onto potato dextrose agar (PDA) medium supplemented with 10 mL/L of a Penicillin-Streptomycin solution (10,000 units penicillin and 10 mg streptomycin/mL, Sigma Aldrich, Saint-Louis, MO, USA) [35]. Fungal isolates were purified on PDA medium and grown for an incubation period of four days at 25°C in the dark followed by three days at room temperature under natural light.

The ratio of the number of tissues contaminated with a species to the total number of tissues analyzed (i.e. the prevalence) was assessed for each species. A chi-squared test was used to determine whether the prevalence of each species on symptomatic tissues differed from that on asymptomatic ones. Pairwise correlations between the prevalence of the six main species were calculated for both symptomatic walnut husks and twigs regardless of the campaign using Pearson’s correlation coefficient (PCC) and plotted using the corrplot R package [59]. Only significant correlations (*p* < 0.05) were plotted. In addition, the frequencies of association at the tissue level were calculated considering only the number of tissues contaminated by the two species involved in the association separately and by the two species together, regardless of the other species. For example, 63 out of 188 symptomatic walnut husks were contaminated by *Neofusicoccum parvum*, and of these, only 21 were contaminated by only this species (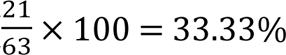 of symptomatic walnut husks contaminated by *Neofusicoccum parvum* only).

A beta-diversity analysis was performed with the species’ prevalence on the symptomatic tissues using a Permutational Multivariate Analysis of Variance (PERMANOVA) followed by a pairwise multilevel comparison using the vegan and pairwise Adonis R packages [60,61].

### Molecular characterization

#### DNA extraction

Mycelium of fungal cultures representative of Botryosphaeriaceae, *Colletotrichum, Diaporthe* and *Fusarium* species (**S1 Table**) was scraped directly from the medium using a sterile scalpel. Extractions were performed using FastDNA™ SpinKit (MP Biomedicals, Fisher Scientific, Waltham, MA, USA) following the manufacturer’s instructions, and DNA solutions were quantified with a NanoDrop 1000 Spectrophotometer (Thermo Fisher Scientific, Waltham, MA, USA).

#### PCR analysis and sequencing

A first molecular characterization was performed by amplifying the internal transcribed spacer (ITS) region using ITS5/ITS4 primers and following White *et al.* (1990) instructions regarding the reaction mixture (**S5 Table**) [62]. Sequencing of the obtained amplicons was performed using the same primer pair by Eurofins Genomics platform (Cologne, Germany). Contig assembly was performed using DNA Baser software and taxonomic assignment to the closest known relatives was performed by comparing contigs with the GenBank database using the “Basic Local Alignment Search Tool” for nucleotide sequences (BlastN; https://blast.ncbi.nlm.nih.gov/Blast.cgi; accessed on 5 September 2022) [63].

For all selected isolates belonging to Botryosphaeriaceae, *Colletotrichum* and *Fusarium*, in-depth molecular characterization was performed using two additional loci for Botryosphaeriaceae (β-tubulin (TUB) and translation elongation factor 1-alpha (EF1-ɑ)) and *Diaporthe* (TUB, and histone H3 (HIS3)), three additional loci for *Fusarium* (EF1-ɑ, RNA polymerase largest subunit (RPB1), and RNA polymerase second largest subunit (RPB2)), and five additional loci for *Colletotrichum* species (TUB, HIS3, glyceraldehyde-3-phosphate dehydrogenase (GAPDH), actin (ACT) and chitin synthase 1 (CHS-1); **S5 Table**). Cycling parameters and reaction mixtures are detailed in **S5 Table**. The PCR products sequencing and contigs assembly were performed as described above. The sequences generated and used in the phylogenetic analysis were deposited in GenBank under accession numbers detailed in **S1 Table**.

#### Phylogenetic analyses

Four separate phylogenetic analyses were conducted to identify Botryosphaeriaceae, *Colletotrichum*, *Diaporthe*, and *Fusarium* species. Reference sequences of targeted loci of closely-related species were retrieved from GenBank (**S1 Table**) to build datasets for each locus and each taxonomic group. Obtained sequences were added to the corresponding dataset and were aligned independently using MAFFT v7 online service [64,65] with the Auto algorithm. The resulting alignments were edited using Gblocks 0.91b [66,67] with all options for a less stringent selection enabled. The individual alignments of the loci were concatenated per taxonomic group as no phylogenetic incongruence was detected (*p =* 0.05 for Botryosphaeriaceae, *p* = 0.15 for *Colletotrichum*, *p* = 0.29 for *Diaporthe,* and *p* = 0.10 for *Fusarium*) by performing an Incongruence Length Difference (ILD) test on 1,000 replicates using PAUP v4.0a166 [68].

Phylogenetic analyses were based on Maximum Likelihood (ML) and Bayesian Inference (BI). For ML analyses, determination of the best models and phylogenetic reconstructions evaluated by 1,000 bootstrap replications were conducted using MEGA-X [69]. For BI analysis, the best models were estimated using the bModelTest add-on in Bayesian Evolutionary Analysis Utility (BEAUti2) [70] simultaneously with the BI analysis using Bayesian Evolutionary Analysis Sampling Trees (BEAST2) v2.6.1 package [71]. The analyses were performed using the Markov Chain Monte Carlo (MCMC) method by performing three independent repetitions of 10,000,000 generations each for each taxonomic group, with sampling a tree every 1,000 generations. Convergence of the independent BI analyses was checked using Tracer v1.7.1 [72], and a consensus tree was obtained for each taxonomic group by a frequency of resampling of 10,000 and a burn-in fixed at 10% using LogCombiner and Treeannotator. *Neoscytalidium dimidiatum* CBS 499.66, *Colletotrichum gloeosporioides* CBS 112999, *Diaporthella corylina* CBS 121124, and *Bisifusarium allantoides* UBOCC-A-120035 were used as outgroups for Botryosphaeriaceae, *Colletotrichum, Diaporthe* and *Fusarium* analyses, respectively. The two phylogenetic trees obtained with ML and BI analyses were compared by each taxonomic group using FigTree v1.4.4 [73]. A tree based on the topology of the ML tree with nodes with bootstrap support < 60% collapsed using TreeCollapserCL 4 [74] was built for each taxonomic group and annotated with bootstrap and posterior probabilities node supports.

### ITS2 metabarcoding analyses

#### DNA extraction of environmental samples

For each collection campaign and for each of the 12 orchards, all samples with the same symptomatology were pooled before being lyophilized and ground as described in Belair *et al.* (2023) [35], yielding a total of 282 samples (94 environmental samples × 3 replicates). Total DNA extractions were performed in triplicate using FastDNA™ Spin Kit and following manufacturer’s instructions. DNA solutions were quantified using a NanoDrop 1000 Spectrophotometer (Thermo Fisher Scientific, Waltham, MA, USA). Extraction blanks were prepared alongside the samples to check for potential contamination.

#### Metabarcoding sequencing and bioinformatic analyses

To study fungal communities, primers ITS3 and ITS4_KYO1 were selected to amplify the ITS2 region [62,75]. This primer pair was previously evaluated as the most appropriate for analyzing fungal communities associated with walnut samples due to its taxonomic resolution and ability to amplify a wide range of fungal diversity [35]. Amplicon libraries and Illumina MiSeq PE 300 bp sequencing were performed in the same run at the McGill University and Génome Québec Innovation Center (Montréal, Canada). Adapter and amplification conditions were the same as described in Belair *et al.* (2023) [35].

Similarly, the workflow used for sequence analysis in this study was the one defined as the most appropriate for the analysis of fungal communities in walnut samples using this primer pair in Belair *et al.* (2023) [35].

#### Alpha- and beta-diversity analysis

The calculation of three alpha-diversity indices (Observed richness, i.e. number of ASVs, Shannon and Simpson’s inverse) for all samples was based on a rarefied dataset at the smallest read counts (540 reads) using the phyloseq R package [76]. These indices were further compared within and between samples by performing an ANOVA and independent sample t.test analyses with the *stat_compare_means* function of the ggpubr R package [77]. In all cases, data were tested for equality and homogeneity of variances.

Relative abundances of taxa across samples were compared after a step of normalization of the non-rarefied dataset using the DESeq2 method [78] as suggested by McMurdie *et al.* (2014) [79]. Differences were expressed as log2 fold changes and their significance levels were tested using Wald test with an alpha threshold set at 0.05 [78]. The normalized datasets were used to produce distance matrices based on Bray-Curtis dissimilarity distance [80] to perform Principal Coordinates Analysis (PCoA) using ggplot2 R package [81]. Statistical analyses were performed with a PERMANOVA followed by a pairwise multilevel comparison using the vegan and pairwiseAdonis R packages [60,61].

Significant correlations between the relative abundance of the top 15 taxa were established for each tissue type using Sparse Correlations for Compositional data algorithm implemented in SparCC python module [82,83]. The co-occurrences networks were plotted for each symptomatic tissue, regardless of the campaign, using the qgraph R package [84]. Only correlations with a correlation strength absolute value greater than 0.3 (|coefficient correlation (=corr)| > 0.3) and p-value inferior to 0.05 were plotted.

### Pathogenicity tests

#### Fungal isolates

Twenty-nine isolates representative of Botryosphaeriaceae (n = 6 *B. dothidea* and n = 6 *N. parvum*), *Diaporthe eres* (n = 6)*, Colletotrichum* (n = 6 *C. godetiae* and n = 6 *C. godetiae)* and *Fusarium juglandicola* (n = 5) were selected for pathogenicity tests (**S1 Table**). All isolates were grown on PDA at 25°C in darkness. Inoculations were carried out using mycelium plugs obtained from the margins of actively growing colonies at least 2 days old.

#### Inoculation of detached twigs

One-year-old detached twigs of the walnut Franquette variety were collected in February 2023 from an experimental orchard belonging to the walnut experimental station (SENuRA) located in Chatte (Isère department, Auvergne-Rhône-Alpes region, France). Twigs approximately 1 m long were stored at 4°C up to 15 days before being processed. To carry out the tests on fresh wood only, the two ends were cut off (about 10 cm) to remove any potentially dried parts. Then, segments of approximately 20 cm including two nodes were cut. Prior to inoculation, the bark surface was disinfected with a 70% ethanol solution. A 6-mm-diameter drill bit was used to drill segments down to the pith. Subsequently, they were inoculated with a 6-mm-diameter mycelium plug which was deposited in contact with the pith, and the inoculated point was immediately covered with petroleum jelly and wrapped with Parafilm. Both ends were sealed with melted paraffin to prevent desiccation and segments were placed in humidity chambers (plastic containers, 27 × 20.5 × 7 cm) filled with sterile filter paper (Grosseron, Couëron, France) and 90 mL sterile distilled water. A total of thirteen segments were placed in each humidity chamber, i.e. ten segments inoculated with one isolate and three control segments including a drilled segment without a PDA plug and petroleum jelly (D), a drilled segment without a PDA plug and covered with petroleum jelly (DJ), and a drilled segment inoculated with a non-inoculated PDA plug and covered with petroleum jelly (DPJ). The humidity chambers were incubated in a climate chamber (Fitotron^➓^ SGC 120, Weiss Technik France S.A.S., Eragny, France) at 25°C with a constant humidity level of 40% and in darkness. The development of the lesion was checked every 10 days in one twig segment and was recorded for all segments when the lesion was almost as long as the segment in case of rapid development (10 days for *B. dothidea*, 11 days for *N. parvum* and 12 days for *D. eres*), or after 31 days in case of slow development (*Colletotrichum* species and *F. juglandicola*). The size of the necrotic lesion in inoculated and control samples was measured after removing the outer bark along the entire length of the segment with a sterile scalpel. The rate of lesion development per day was then calculated (mm/day). In order to validate the Koch postulate, the margins of the lesion or inoculated points when no lesion were detected, as well as control twigs, were cut and placed onto PDA supplemented with the same antibiotic solution as described in **Collection of fungal isolates** and incubated at 25°C in the dark as described above. The reisolation rate for each isolate was then calculated as the number of positive inoculated points to the total number of inoculated points.

#### Inoculations of detached walnut fruits

Green fruits of the walnut variety Fernor were collected in September 2023 from an experimental orchard belonging to the SENuRA. Fruits were first washed with sterile distilled water, then surface disinfected in a 70% ethanol solution bath for 1 min and finally rinsed in two sterile distilled water for 1 min each and dried on sterile paper sheets. A 3 to 6 mm diameter wound was made in the center of each fruit using a 2 mL pipette tip and was inoculated by placing a mycelium plug of the same diameters. Inoculation points were then covered with petroleum jelly. Isolates used for these pathogenicity tests are listed in **S1 Table**. Inoculated fruits were placed in humidity chambers (plastic containers, 22.5 × 14.5 × 4.5 cm) filled with sterile filter paper and 4 mL sterile distilled water the first day and 8 mL sterile distilled water the following days. Humidity chambers contained 10 green fruits inoculated with one isolate and were then incubated in a climate chamber (Memmert ICH260L, Memmert GmbH, Schwabach, Germany) at 25°C with a constant humidity of 80% and a 16 hours photoperiod. Among these, three humidity chambers contained 10 fruits each representing control conditions, i.e. non-drilled green fruits (ND), non-inoculated drilled green fruits covered with petroleum jelly (DJ), and drilled green fruits inoculated with non-inoculated PDA plug and covered with petroleum jelly (DPJ). Disease severity was frequently assessed until fruits with 90% to 100% of their surface affected were observed, i.e. 12 days after inoculation. The average of the major and minor diameters of each lesion was calculated for each fruit and isolate in order to assess the area of the necrotic lesion and the percentage of affected fruit surface. The mean relative area under the disease progress curve (RAUDPC) was obtained by trapezoidal integration for each fruit, using the agricolae R package [85,86]. In order to fulfill Koch’s postulates, fungal reisolation was performed from the margins (three pieces) of the necrotic lesions for both inoculated green fruits and controls. All the reisolations were performed onto PDA supplemented with 0.1 g/L of tetracyclin hydrochloride CELLPURE^➓^

(Carl Roth, Karlsruhe, Germany) and incubated at 25°C with a 16 hours photoperiod. The reisolation rate for each isolate was then calculated as the number of positive necrotic inoculated points to the total number of necrotic inoculated points.

#### Inoculations of attached twigs under field conditions

One-year-old twigs of the walnut Franquette variety were selected from an experimental orchard belonging to the walnut experimental station located in Creysse (Lot department, Occitanie region, France). The area of inoculation was surface-disinfected with a 70% ethanol solution. As described in **Inoculations of detached twigs**, twigs were drilled to the pith using a 6-mm-diameter drill bit and inoculations were performed using mycelium plug covered with petroleum jelly and Parafilm (isolates are listed in **S1 Table**). Two walnut trees were selected, and twigs were randomly inoculated in order to obtain 13 inoculated twigs per isolate. Additionally, 26 twigs represented control conditions: 13 twigs were drilled and inoculated with a non-inoculated PDA plug (DPJ) and 13 others were not drilled (ND). All were wrapped with Parafilm. Lesion lengths were recorded 6 months after inoculation. All inoculated and control twigs were used for reisolations, which were performed by plating the margins of necrosis or wounded point onto PDA supplemented with 0.5 g/L of tetracyclin hydrochloride (Thermo Fisher Scientific, Waltham, MA, USA), and incubated at ambient temperature and luminosity. The reisolation rate for each isolate was then calculated as the number of positive inoculated points to the total number of inoculated points.

#### Data analysis

In all cases, mean lesion length and RAUDPC were compared according to Kruskal-Wallis and Dunn’ test at ɑ = 0.05. Statistical comparisons were performed between all tested species and controls (regardless of isolates) and then between all isolates of a species and controls.

### Meteorological data

The meteorological data were retrieved from Meteo France [87]. The mean daily temperature and rainfall were collected from April to August 2020, 2021 and 2022, from the six weather stations closest to the studied orchards (**S3 Table**). Three stations were located in the Auvergne-Rhône-Alpes region, in the South-East production area, while the three others were located in the Nouvelle-Aquitaine region, in the South-West production area. The data were statistically compared using ANOVA followed by Tukey’s HSD test (ɑ = 0.05). Correlations between the prevalence of the main species and weather variables were tested by performing Canonical Correspondence Analysis (CCA) using the vegan R package [61].

## Acknowledgements

We would like to thank the growers for giving us access to their orchards in this project.

## Data Availability

The metabarcoding data presented in this study have been deposited at the NCBI and are openly available under the Bioproject SUB15096080.

## Author Contributions

FP and AP acquired funding and supervised the scientific management of the PhD work, along with GLF. CM, M-NH, AM and YL sampled and supplied walnut husks and twigs. AM, YL, LM, MB, AP and FP co-developed the protocol for pathogenicity tests on detached twigs. CM and M-NH performed pathogenicity tests on detached walnut fruits and in walnut orchards (with the technical support of MD), respectively. BR and MB analyzed meteorological data. AH-S and MB performed multilocus amplifications experiments, and MB performed phylogenetic analyses. LM and MB performed and analyzed pathogenicity tests on detached walnut twigs. ST provides crucial technical support. MB conducted culture-dependent and - independent experiments and analyses, and generated all the tables and figures. MB, AP, and FP wrote the manuscript. All authors have read and approved the manuscript.

## Supporting information

**S1 Fig. Maximum Likelihood phylogenetic consensus tree of fungal species belonging to Botryosphaeriaceae based on the concatenation of the sequences of three genes (ITS, EF1- ɑ and TUB).** Maximum Likelihood bootstrap supports (BS; > 60%) and Bayesian posterior probabilities (PP) are represented at nodes (BS/PP). *Neoscytalidium dimidiatum* CBS 499.66 was used as an outgroup.

**S2 Fig. Maximum Likelihood phylogenetic consensus tree of fungal species belonging to *Colletotrichum* based on the concatenation of the sequences of six genes (ITS, ACT, CHS-1, GAPDH, HIS3 and TUB).** Maximum Likelihood bootstrap supports (BS; > 60%) and Bayesian posterior probabilities (PP) are represented at nodes (BS/PP). *Colletotrichum gloeosporioides* CBS 112999 was used as an outgroup.

**S3 Fig. Maximum Likelihood phylogenetic consensus tree of fungal species belonging to *Diaporthe* based on the concatenation of the sequences of two genes (HIS3 and TUB).** Maximum Likelihood bootstrap supports (BS; > 60%) and Bayesian posterior probabilities (PP) are represented at nodes (BS/PP). *Diaporthella corylina* CBS 121124 was used as an outgroup.

**S4 Fig. Maximum Likelihood phylogenetic consensus tree of fungal species belonging to *Fusarium* based on the concatenation of the sequences of four genes (ITS, EF1-ɑ, RPB1 and RPB2).** Maximum Likelihood bootstrap supports (BS; > 60%) and Bayesian posterior probabilities (PP) are represented at nodes (BS/PP). *Bisifusarium allantoides* UBOCC-A-120035 was used as an outgroup.

**S5 Fig. Summary of the meteorological data (rainfall (mm) and temperature (°C)) from the meteorological station of the South-Eats production area.** (A) CHATTE_SAPC, (B) SERRE-NERPOL_SAPC, and (C) ST-JEAN-EN-ROYANS. The meteorological data are represented from April to August in 2020, 2021 and 2022.

**S6 Fig. Summary of the meteorological data (rainfall (mm) and temperature (°C)) from the meteorological station of the South-West production area.** (A) LES-EYZIES-DE-TAYAC, (B) MONPAZIER, and (C) SALIGNAC-EYVIGUES. The meteorological data are represented from April to August in 2020, 2021 and 2022.

**S1 Table. Fungal isolates used in the phylogenetic analysis and their corresponding GenBank accession numbers.**

**S2 Table. Frequency of reisolation (%) of fungal isolates from 1-year-old detached and attached twigs, and detached green fruits of English walnut used in pathogenicity tests.**

**S3 Table. Description and location of weather stations used in this study.**

**S4 Table. Mean temperature (°C) and mean cumulative rainfall (mm) for each month from April to August in 2020, 2021 and 2022 in both production areas.**

**S5 Table. Primers, cycling parameters and reaction mixtures used for molecular characterization.**

## References

1. Martínez-García PJ, Crepeau MW, Puiu D, Gonzalez-Ibeas D, Whalen J, Stevens KA, et al. The walnut (*Juglans regia*) genome sequence reveals diversity in genes coding for the biosynthesis of non-structural polyphenols. Plant J. 2016;87: 507–532. doi:10.1111/tpj.13207

2. FAOSTAT. Statistical database - World and France total production. Walnut with shell. License: CC BY-NC-SA 3.0 IGO. 2021 [cited 1 Feb 2023]. Available: https://www.fao.org/faostat/en/#data/QCL

3. Verhaeghe A. Anthracnose - Gnomonia leptostyla. 2014 Apr. Report No.: 1. Available: https://senura.com/images/DOCUMENTS/DOCUMENTS_TECHNIQUES/Plaquette_ANTHRACNOSE_2014.pdf

4. Radix P, Seigle-Murandi F, Charlot G. Walnut blight: development of fruit infection in two orchards. Crop Prot. 1994;13: 629–631. doi:10.1016/0261-2194(94)90010-8

5. Da Lio D, Cobo-Díaz JF, Masson C, Chalopin M, Kebe D, Giraud M, et al. Combined Metabarcoding and Multi-locus approach for Genetic characterization of *Colletotrichum* species associated with common walnut (*Juglans regia*) anthracnose in France. Sci Rep. 2018;8: 10765. doi:10.1038/s41598-018-29027-z

6. Laloum Y, Verhaeghe A, Moronvalle A, Masson C, Hebrard M-N, Picot A, et al. Projet CARIBOU : Élancer la recherche pour cerner le dépérissement du noyer. INFOS CTIFL. 2022;386: 48–58.

7. Trouillas FP, Hébrard M-N, Aurelle V, Jargeat P, Tranchand E. Etiology of walnut fruit rot, spur and twig dieback in southwestern France. Acta Hortic. 2023. Forthcoming.

8. Chen SF, Morgan DP, Hasey JK, Anderson K, Michailides TJ. Phylogeny, morphology, distribution, and pathogenicity of Botryosphaeriaceae and Diaporthaceae from English walnut in California. Plant Dis. 2014;98: 636–652. doi:10.1094/PDIS-07-13-0706-RE

9. Eichmeier A, Pecenka J, Spetik M, Necas T, Ondrasek I, Armengol J, et al. Fungal Trunk Pathogens Associated With *Juglans regia* in the Czech Republic. Plant Dis. 2020;104: 761–771. doi:10.1094/PDIS-06-19-1308-RE

10. López-Moral A, Lovera M, del Carmen Raya M del C, Cortés-Cosano N, Arquero O, Trapero A, et al. Etiology of Branch Dieback and shoot blight of English walnut caused by Botryosphaeriaceae and *Diaporthe* species in southern Spain. Plant Dis. 2020;104: 533–550. doi:10.1094/PDIS-03-19-0545-RE

11. Gusella G, Giambra S, Conigliaro G, Burruano S, Polizzi G. Botryosphaeriaceae species causing canker and dieback of English walnut (*Juglans regia*) in Italy. For Path. 2021;51: e12661. doi:10.1111/efp.12661

12. Mihaescu C, Dunea D, Bășa AG, Frasin LN. Characteristics of *Phomopsis juglandina* (Sacc.) Hohn. Associated with Dieback of Walnut in the Climatic Conditions of Southern Romania. Agronomy. 2021;11: 46. doi:10.3390/agronomy11010046

13. Zabiák A, Kovács C, Takács F, Pál K, Peles F, Fekete E, et al. *Diaporthe* and *Diplodia* Species Associated with Walnut (*Juglans regia* L.) in Hungarian Orchards. Horticulturae. 2023;9: 205. doi:10.3390/horticulturae9020205

14. Hilário S, Gonçalves MFM. Mechanisms Underlying the Pathogenic and Endophytic Lifestyles in *Diaporthe*: An Omics-Based Approach. Horticulturae. 2023;9: 423. doi:10.3390/horticulturae9040423

15. Slippers B, Wingfield MJ. Botryosphaeriaceae as endophytes and latent pathogens of woody plants: diversity, ecology and impact. Fungal Biol Rev. 2007;21: 90–106. doi:10.1016/j.fbr.2007.06.002

16. Guerin-Dubrana L, Fontaine F, Mugnai L. Grapevine trunk disease in European and Mediterranean vineyards: occurrence, distribution and associated disease-affecting cultural factors. Phytopathol Mediterr. 2019;58: 49–71. doi:10.14601/Phytopathol_Mediterr-25153

17. López-Moral A, del Carmen Raya M, Ruiz-Blancas C, Medialdea I, Lovera M, Arquero O, et al. Aetiology of branch dieback, panicle and shoot blight of pistachio associated with fungal trunk pathogens in southern Spain. Plant Pathol. 2020;69: 1237–1269. doi:10.1111/ppa.13209

18. Batista E, Lopes A, Alves A. Botryosphaeriaceae species on forest trees in Portugal: diversity, distribution and pathogenicity. Eur J Plant Pathol. 2020;158: 693–720. doi:10.1007/s10658-020-02112-8

19. Moricca S, Innocenti G, Ragazzi A. Epidemiological Investigations Shed Light on the Ecological Role of the Endophyte *Phomopsis quercina* in Mediterranean Oak Forests. In: Pirttilä AM, Frank AC, editors. Endophytes of Forest Trees. Cham: Springer International Publishing; 2018. pp. 207–227. doi:10.1007/978-3-319-89833-9_10

20. Agustí-Brisach C, Moral J, Felts D, Trapero A, Michailides TJ. Interaction Between *Diaporthe rhusicola* and *Neofusicoccum mediterraneum* Causing Branch Dieback and Fruit Blight of English Walnut in California, and the Effect of Pruning Wounds on the Infection. Plant Dis. 2019;103: 1196–1205. doi:10.1094/PDIS-07-18-1118-RE

21. Wallis CM, Gorman Z, Galarneau ER-A, Baumgartner K. Mixed infections of fungal trunk pathogens and induced systemic phenolic compound production in grapevines. Front Fungal Biol. 2022;3. doi:10.3389/ffunb.2022.1001143

22. Antony S, Billones-Baaijens R, Stodart BJ, Steel CC, Lang MD, Savocchia S. Incidence and distribution of Botryosphaeriaceae species associated with dieback in walnut orchards in Australia. Plant Pathol. 2023;72: 610–622. doi:10.1111/ppa.13685

23. Jiménez Luna I, Besoain X, Saa S, Peach-Fine E, Cadiz Morales F, Riquelme N, et al. Identity and pathogenicity of Botryosphaeriaceae and Diaporthaceae from *Juglans regia* in Chile. Phytopathol Mediterr. 2022;61: 79–94. doi:10.36253/phyto-12832

24. Li G, Liu F, Li J, Liu Q, Chen S. Characterization of *Botryosphaeria dothidea* and *Lasiodiplodia pseudotheobromae* from English Walnut in China. J Phytopathol. 2016;164: 348–353. doi:10.1111/jph.12422

25. Cannon PF, Damm U, Johnston PR, Weir BS. *Colletotrichum* – current status and future directions. Stud Mycol. 2012;73: 181–213. doi:10.3114/sim0014

26. Belisario A, Santori A, Potente G, Fiorin A, Saphy B, Reigne JL, et al. Brown apical necrosis (BAN): a fungal disease causing fruit drop of English walnut. Acta Hortic. 2010;861: 449–452. doi:10.17660/ActaHortic.2010.861.63

27. Wang YX, Chen JY, Xu XW, Li DW, Wang QZ. First Report of Brown Apical Necrosis of Walnut Fruit Caused by *Fusarium avenaceum* in Hubei, China. Plant Dis. 2019;103: 2956. doi:10.1094/PDIS-05-19-0954-PDN

28. Wang S, Tan Y, Li S, Zhu T. Structural and Dynamic Analysis of Leaf-Associated Fungal Community of Walnut Leaves Infected by Leaf Spot Disease Based Illumina High-Throughput Sequencing Technology. Pol J Microbiol. 2022;71: 429–441. doi:10.33073/pjm-2022-038

29. Wang X, Li K, Han M, Zhang W, Li X, Ma D, et al. Isolation and identification of endophytic fungi in walnut. IOP Conf Ser: Earth Environ Sci. 2020;508: 012138. doi:10.1088/1755-1315/508/1/012138

30. Moragrega C, Özaktan H. Apical Necrosis of Persian (english) Walnut (*Juglans regia*): An Update. J Plant Pathol. 2010;92: S67–S71. doi:https://www.jstor.org/stable/41998757

31. Chen W, Swart WJ. First Report of Stem Canker of English Walnut Caused by *Fusarium solani* in South Africa. Plant Dis. 2000;84: 592–592. doi:10.1094/PDIS.2000.84.5.592A

32. Mulero-Aparicio A, Agustí-Brisach C, Raya M del C, Lovera M, Arquero O, Trapero A. First Report of *Fusarium solani* Causing Stem Canker in English Walnut in Spain. Plant Dis. 2019;103: 3281–3281. doi:10.1094/PDIS-06-19-1163-PDN

33. Polat Z, Gültekin MA, Palacıoğlu G, Bayraktar H, Özer N, Yılmaz S. First report of *Neocosmospora solani* causing stem canker on *Juglans regia* in Turkey. J Plant Pathol. 2020;102: 1289–1289. doi:10.1007/s42161-020-00570-x

34. Tuerdi M, Lv C, Dan H, Shan L, Zhang R, Chen X. First Report of *Fusarium solani* Associated with Twig Canker of English Walnut (*Juglans regia*) in Xinjiang, China. Plant Dis. 2023;107: 2858. doi:10.1094/PDIS-03-23-0430-PDN

35. Belair M, Pensec F, Jany J-L, Floch GL, Picot A. Profiling walnut fungal pathobiome associated with walnut dieback using community-targeted DNA metabarcoding. Plants. 2023;12: 2383. doi:10.3390/plants12122383

36. Michailides TJ, Chen S, Morgan D, Felts D, Nouri MT, Puckett R, et al. Managing Botryosphaeria/Phomopsis cankers and anthracnose blight of walnut in California. Walnut Research Reports. California Walnut Board, Folsom, CA; 2013. pp. 325–346.

37. Belair M, Picot A, Lepais O, Masson C, Hébrard M-N, Moronvalle A, et al. Genetic diversity and population structure of <i>Botryosphaeria dothidea</i and *Neofusicoccum parvum* on English walnut (*Juglans regia* L.) in France. Sci Rep. 2024;14: 19817. doi:10.1038/s41598-024-67613-6

38. Yildiz A, Benlioglu S, Benlioglu K, Korkom Y. Occurrence of twig blight and branch dieback of walnut caused by Botryosphaeriaceae species in Turkey. J Plant Dis Protect. 2022;129: 687–693. doi:10.1007/s41348-022-00591-x

39. Novak A, Ivić D, Sever Z, Fazinić T, Šimunac K. Botryosphaeria canker of walnut in Croatia. Glasilo biljne zaštite. 2018;18: 316–321. doi:https://hrcak.srce.hr/236900

40. Varjas V, Lakatos T, Tóth T, Kovács C. First Report of *Colletotrichum godetiae* Causing Anthracnose and Twig Blight on Persian Walnut in Hungary. Plant Dis. 2020;105: 702– 702. doi:10.1094/PDIS-03-20-0607-PDN

41. Crous PW, Cowan DA, Maggs-Kölling G, Yilmaz N, Thangavel R, Wingfield MJ, et al. Fungal Planet description sheets: 1182–1283. Persoonia - Molecular Phylogeny and Evolution of Fungi. 2021;46: 313–528. doi:10.3767/persoonia.2021.46.11

42. Estrada AB, Dodd JC, Jeffries P. Effect of humidity and temperature on conidial germination and appressorium development of two Philippine isolates of the mango anthracnose pathogen *Colletotrichum gloeosporioides*. Plant Pathol. 2000;49: 608–618. doi:10.1046/j.1365-3059.2000.00492.x

43. Hall BH, Pederick SJ, McKay SF. Understanding infection of pistachio by *Colletotrichum acutatum*. Acta Hortic. 2016;1109: 215–222. doi:10.17660/ActaHortic.2016.1109.35

44. Kamle M, Kumar P. *Colletotrichum gloeosporioides: Pathogen of Anthracnose Disease in Mango (*Mangifera indica* L.)*. In: Kumar P, Gupta VK, Tiwari AK, Kamle M, editors. Current Trends in Plant Disease Diagnostics and Management Practices. Cham: Springer International Publishing; 2016. pp. 207–219. doi:10.1007/978-3-319-27312-9_9

45. Salotti I, Liang Y-J, Ji T, Rossi V. Development of a model for Colletotrichum diseases with calibration for phylogenetic clades on different host plants. Front Plant Sci. 2023;14. doi:10.3389/fpls.2023.1069092

46. González-Domínguez E, Caffi T, Languasco L, Latinovic N, Latinovic J, Rossi V. Dynamics of *Diaporthe ampelina* Conidia Released from Grape Canes that Overwintered in the Vineyard. Plant Dis. 2021;105: 3092–3100. doi:10.1094/PDIS-12-20-2639-RE

47. Moral J, Morgan D, Trapero A, Michailides TJ. Ecology and Epidemiology of Diseases of Nut Crops and Olives Caused by Botryosphaeriaceae Fungi in California and Spain. Plant Dis. 2019;103: 1809–1827. doi:10.1094/PDIS-03-19-0622-FE

48. Arciuolo R, Camardo Leggieri M, Chiusa G, Castello G, Genova G, Spigolon N, et al. Ecology of *Diaporthe eres*, the causal agent of hazelnut defects. PLOS ONE. 2021;16: e0247563. doi:10.1371/journal.pone.0247563

49. Salotti I, Ji T, Rossi V. Temperature requirements of *Colletotrichum* spp. belonging to different clades. Front Plant Sci. 2022;13. doi:10.3389/fpls.2022.953760

50. Penet L, Guyader S, Pétro D, Salles M, Bussière F. Direct Splash Dispersal Prevails over Indirect and Subsequent Spread during Rains in *Colletotrichum gloeosporioides* Infecting Yams. PLOS ONE. 2014;9: e115757. doi:10.1371/journal.pone.0115757

51. Velásquez AC, Castroverde CDM, He SY. Plant and pathogen warfare under changing climate conditions. Curr Biol. 2018;28: R619–R634. doi:10.1016/j.cub.2018.03.054

52. Zhao J, Lu Z, Wang L, Jin B. Plant Responses to Heat Stress: Physiology, Transcription, Noncoding RNAs, and Epigenetics. Int J Mol Sci. 2020;22: 117. doi:10.3390/ijms22010117

53. Iqbal M, Broberg A, Andreasson E, Stenberg JA. Biocontrol potential of beneficial fungus *Aureobasidium pullulans* against *Botrytis cinerea* and *Colletotrichum acutatum*. Phytopathology. 2023;113: 1428–1438. doi:10.1094/PHYTO-02-23-0067-R

54. Pinto C, Custódio V, Nunes M, Songy A, Rabenoelina F, Courteaux B, et al. Understand the Potential Role of *Aureobasidium pullulans*, a Resident Microorganism From Grapevine, to Prevent the Infection Caused by *Diplodia seriata*. Front Microbiol. 2018;9: 3047. doi:10.3389/fmicb.2018.03047

55. Rueda-Mejia MP, Nägeli L, Lutz S, Hayes RD, Varadarajan AR, Grigoriev IV, et al. Genome, transcriptome and secretome analyses of the antagonistic, yeast-like fungus *Aureobasidium pullulans* to identify potential biocontrol genes. Microb Cell. 2021;8: 184–202. doi:10.15698/mic2021.08.757

56. Nerva L, Garcia JF, Favaretto F, Giudice G, Moffa L, Sandrini M, et al. The hidden world within plants: metatranscriptomics unveils the complexity of wood microbiomes. J Exp Bot. 2022;73: 2682–2697. doi:10.1093/jxb/erac032

57. Cai F, Yang C, Ma T, Jin M, Cui L. First report of *Boeremia exigua* var. *exigua* causing branch blight on walnut in China. Plant Dis. 2021;105: 3291. doi:10.1094/PDIS-02-21-0382-PDN

58. Wang F, Dun C, Tang T, Duan Y, Guo X, You J. *Boeremia exigua* Causes Leaf Spot of Walnut Trees (*Juglans regia*) in China. Plant Dis. 2022;106: 1993. doi:10.1094/PDIS-10-21-2304-PDN

59. Wei T, Simko V. corrplot: A visual exploratory tool on correlation matrix. 2021. Available: https://github.com/taiyun/corrplot

60. Arbizu PM. pairwiseAdonis: Pairwise Multilevel Comparison using Adonis. 2023. Available: https://github.com/pmartinezarbizu/pairwiseAdonis

61. Oksanen J, Simpson GL, Blanchet FG, Kindt R, Legendre P, Minchin PR, et al. vegan: Community Ecology Package. 2022. Available: https://cran.r-project.org/web/packages/vegan/index.html

62. White TJ, Bruns T, Lee S, Taylor J. Amplification and direct sequencing of fungal ribosomal RNA genes for phylogenetics. Academic press. PCR protocols: a guide to methods and amplifications. Academic press. Cambridge, MA, USA: Academic Press; 1990. pp. 315–322. doi:10.1016/B978-0-12-372180-8.50042-1

63. Altschul SF, Gish W, Miller W, Myers EW, Lipman DJ. Basic local alignment search tool. J Mol Biol. 1990;215: 403–410. doi:10.1016/S0022-2836(05)80360-2

64. Katoh K, Rozewicki J, Yamada KD. MAFFT online service: multiple sequence alignment, interactive sequence choice and visualization. Brief Bioinform. 2019;20: 1160–1166. doi:10.1093/bib/bbx108

65. Kuraku S, Zmasek CM, Nishimura O, Katoh K. aLeaves facilitates on-demand exploration of metazoan gene family trees on MAFFT sequence alignment server with enhanced interactivity. Nucleic Acids Res. 2013;41: W22–W28. doi:10.1093/nar/gkt389

66. Castresana J. Selection of Conserved Blocks from Multiple Alignments for Their Use in Phylogenetic Analysis. Mol Biol Evol. 2000;17: 540–552. doi:10.1093/oxfordjournals.molbev.a026334

67. Talavera G, Castresana J. Improvement of Phylogenies after Removing Divergent and Ambiguously Aligned Blocks from Protein Sequence Alignments. Syst Biol. 2007;56: 564–577. doi:10.1080/10635150701472164

68. Swofford D. PAUP*. Phylogenetic analysis using parsimony and other methods. Version 4.0. Sinauer Associates. Sunderland, Massachusetts; 2003.

69. Kumar S, Stecher G, Li M, Knyaz C, Tamura K. MEGA X: Molecular Evolutionary Genetics Analysis across Computing Platforms. Mol Biol Evol. 2018;35: 1547–1549. doi:10.1093/molbev/msy096

70. Bouckaert RR, Drummond AJ. bModelTest: Bayesian phylogenetic site model averaging and model comparison. BMC Evol Biol. 2017;17: 42. doi:10.1186/s12862-017-0890-6

71. Bouckaert R, Vaughan TG, Barido-Sottani J, Duchêne S, Fourment M, Gavryushkina A, et al. BEAST 2.5: An advanced software platform for Bayesian evolutionary analysis. PLOS Comput Biol. 2019;15: e1006650. doi:10.1371/journal.pcbi.1006650

72. Rambaut A, Drummond AJ, Xie D, Baele G, Suchard MA. Posterior Summarization in Bayesian Phylogenetics Using Tracer 1.7. Syst Biol. 2018;67: 901–904. doi:10.1093/sysbio/syy032

73. Rambaut A. FigTree. 2006 [cited 26 Jun 2024]. Available: http://tree.bio.ed.ac.uk/software/figtree/

74. Hodcroft E. TreeCollapserCL4. 28 Jun 2024 [cited 28 Jun 2024]. Available: http://emmahodcroft.com/TreeCollapseCL.html

75. Toju H, Tanabe AS, Yamamoto S, Sato H. High-Coverage ITS Primers for the DNA-Based Identification of Ascomycetes and Basidiomycetes in Environmental Samples. PLoS ONE. 2012;7: e40863. doi:10.1371/journal.pone.0040863

76. McMurdie PJ, Holmes S. phyloseq: An R Package for Reproducible Interactive Analysis and Graphics of Microbiome Census Data. PLOS ONE. 2013;8: e61217. doi:10.1371/journal.pone.0061217

77. Kassambara A. ggpubr: “ggplot2” Based Publication Ready Plots. 2023. Available: https://cran.r-project.org/web/packages/ggpubr/index.html

78. Love MI, Huber W, Anders S. Moderated estimation of fold change and dispersion for RNA-seq data with DESeq2. Genome Biol. 2014;15: 550. doi:10.1186/s13059-014-0550-8

79. McMurdie PJ, Holmes S. Waste not, want not: why rarefying microbiome data is inadmissible. PLOS Comput Biol. 2014;10: e1003531. doi:10.1371/journal.pcbi.1003531

80. Bray JR, Curtis JT. An Ordination of the Upland Forest Communities of Southern Wisconsin. Ecol Monogr. 1957;27: 326–349. doi:10.2307/1942268

81. Wickham H. ggplot2 : Elegant Graphics for Data Analysis. New-York: Springer International Publishing; 2016. doi:10.1007/978-3-319-24277-4

82. Friedman J, Alm EJ. Inferring Correlation Networks from Genomic Survey Data. PLOS Comput Biol. 2012;8: e1002687. doi:10.1371/journal.pcbi.1002687

83. Legorreta D. SparCC. 2021. Available: https://github.com/dlegor/SparCC

84. Epskamp S, Cramer AOJ, Waldorp LJ, Schmittmann VD, Borsboom D. qgraph: Network Visualizations of Relationships in Psychometric Data. J Stat Softw. 2012;48: 1–18. doi:10.18637/jss.v048.i04

85. de Mendiburu F. agricolae: Statistical Procedures for Agricultural Research. 2023. Available: https://cran.r-project.org/web/packages/agricolae/index.html

86. Campbell CL, Madden LV. Introduction to plant disease epidemiology. Wiley. New York; 1990.

87. Meteo France. METEO-FRANCE : Publithèque. 2023. Available: https://publitheque.meteo.fr/okapi/accueil/okapiWebPubli/index.jsp

